# Evolutionarily ancient BAH-PHD protein mediates Polycomb silencing

**DOI:** 10.1101/868117

**Authors:** Elizabeth T. Wiles, Kevin J. McNaught, Saumya M. De Silva, Gurmeet Kaur, Jeanne M. Selker, Tereza Ormsby, L. Aravind, Catherine A. Musselman, Eric U. Selker

## Abstract

Methylation of histone H3 lysine 27 (H3K27) is widely recognized as a transcriptionally repressive chromatin modification but the mechanism of repression remains unclear. We devised and implemented a forward genetic scheme to identify factors required for H3K27 methylation-mediated silencing in the filamentous fungus *Neurospora crassa* and identified a bromo-adjacent homology (BAH)-plant homeodomain (PHD)-containing protein, EPR-1 (<u>E</u>ffector of <u>P</u>olycomb <u>R</u>epression <u>1</u>; NCU07505). EPR-1 associates with H3K27 methylation *in vivo* and *in vitro*, and loss of EPR-1 de-represses H3K27-methylated genes without loss of H3K27 methylation. EPR-1 is not fungal-specific; orthologs of EPR-1 are present in a diverse array of eukaryotic lineages, suggesting an ancestral EPR-1 was a component of a primitive Polycomb repression pathway.

**Significance:** Polycomb group (PcG) proteins are employed by a wide variety of eukaryotes for the maintenance of gene repression. Polycomb repressive complex 2 (PRC2), a multimeric complex of PcG proteins, catalyzes the methylation of histone H3 lysine 27 (H3K27). In the filamentous fungus, *Neurospora crassa*, H3K27 methylation represses scores of genes, despite the absence of canonical H3K27 methylation effectors that are present in plants and animals. We report the identification and characterization of an H3K27 methylation effector, EPR-1, in *N. crassa* and demonstrate its widespread presence and early eukaryotic origins with phylogenetic analyses. These findings indicate that an ancient EPR-1 may have been part of a nascent Polycomb repression system in eukaryotes.

## Introduction

The establishment and maintenance of transcriptionally repressive chromatin is critical for the development of multicellular organisms (1-4). Polycomb group (PcG) proteins, originally discovered in *Drosophila melanogaster* (5), form multiple complexes that maintain such chromatin repression (6). Although the composition of PcG complexes varies, a few core constituents define two major classes of chromatin-modifying complexes, namely Polycomb repressive complex 1 (PRC1) and Polycomb repressive complex 2 (PRC2) (7). According to the ‘classical model,’ PcG-mediated gene silencing is initiated by targeting of PRC2 to chromatin (8), which catalyzes methylation of H3K27 (9). Canonical PRC1, which contains a chromodomain protein (*e*.*g*. Polycomb in *Drosophila melanogaster* and CBX2/4/6-8 in mammals), recognizes tri-methylated H3K27 (10), catalyzes monoubiquitination of neighboring histone H2A lysine 119 by RING1A/B (11), and promotes chromatin compaction (12, 13). In reality, this hierarchical recruitment model is an oversimplification, as PRC1 can be recruited to PcG targets irrespective of PRC2 activity (14) and PRC1 presence is required for stable PRC2 association at many Polycomb Response Elements in *Drosophila melanogaster* (15). Interdependence of these complexes has limited our understanding of their respective roles and the function of their associated chromatin ‘marks’ on gene repression.

While plants and animals utilize distinct sets of accessory proteins to recognize methylated H3K27 (10, 16-18), they are generally thought to mediate repression in the context of a canonical PRC1 complex (7, 19-21). In fungal lineages that employ H3K27 methylation as a repressive chromatin mark, however, core PRC1 components are notably absent (7). This raises the question of how H3K27 methylation mediates repression in the absence of PRC1. It suggests that either: 1. H3K27 methylation *per se* may be repressive, or 2. There is a ‘reader’ of H3K27 methylation that functions outside the context of canonical PRC1.

To elucidate the repressive mechanism of H3K27 methylation in fungi, we developed and employed a forward genetics approach to identify effectors of Polycomb repression using *Neurospora crassa*. H3K27 methylation covers approximately 7% of the *N. crassa* genome and is responsible for the repression of scores of genes (22, 23). We found four mutant alleles of an undescribed gene (*NCU07505*) that we show is critical for H3K27 methylation-mediated silencing and therefore named it *e*ffector of *P*olycomb *r*epression *1 (epr-1)*. It encodes a protein with a bromo-adjacent homology (BAH) domain and plant homeodomain (PHD) finger. Although *epr-1* mutants display phenotypic and gene expression changes similar to strains lacking PRC2 components, H3K27 methylation is essentially unaffected. We demonstrate that EPR-1 forms nuclear foci, reminiscent of Polycomb bodies (24), and its genomic distribution is limited to, and dependent upon, H3K27-methylated chromatin, which it recognizes directly through its BAH domain. Finally, we discover that EPR-1 orthologs are widely distributed across eukaryotes, contrary to previous reports (21, 25, 26), suggesting an ancient role of EPR-1 homologs in Polycomb repression that was then lost on multiple occasions in certain lineages.

## Results

### Genetic selection for factors necessary for H3K27 methylation-mediated repression

In an effort to identify factors required for H3K27 methylation-mediated repression, we engineered a strain of *N. crassa* in which we replaced the open reading frames of two PRC2-repressed genes (23), *NCU05173* and *NCU07152*, with the antibiotic-resistance genes *hph* and *nat-1*, respectively (Fig. 1a). Strains that bear these gene replacements and lack the H3K27 methyltransferase (SET-7) are resistant to Hygromycin B and Nourseothricin, whereas a wild-type strain with these gene replacements is sensitive to these drugs (Fig. 1c). We subjected conidia collected from such an antibiotic-sensitive, otherwise wild-type strain to ultraviolet (UV) mutagenesis and selected for mutants that derepressed both the *hph* and *nat-1* genes (Fig. 1b). One mutant isolated in this manner and characterized here is *e*ffector of *p*olycomb *r*epression *1* (*epr-1*) (Fig. 1c).

**Fig. 1.**
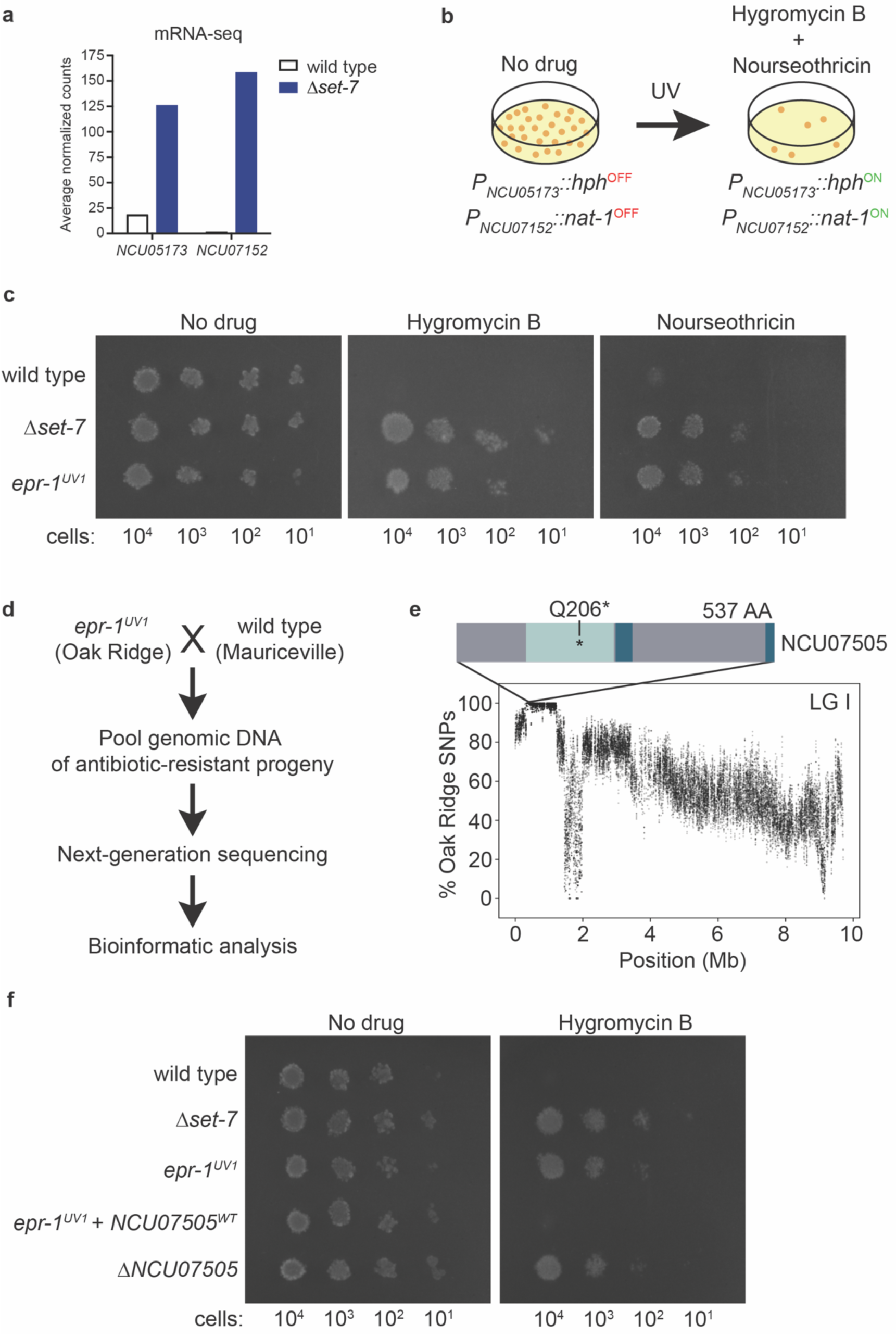
Forward genetics identifies a novel gene, *epr-1*, required for H3K27 methylation-mediated repression. **a**, mRNA-seq results for two genes repressed by the *N. crassa* H3K27 methyltransferase, encoded by *set-7* (23). **b**, Selection scheme, utilizing reporter genes illustrated in **a**, to identify factors required for H3K27 methylation-mediated silencing. **c**, Serial dilution spot test silencing assay for the indicated strains plated on the indicated media. All strains harbor *P*_*NCU05173*_*::hph* and *P*_*NCU07152*_*::nat-1*. **d**, Scheme for genetic mapping of critical mutation in *epr-1*^*UV1*^. **e**, Whole genome sequencing of pooled *epr-1*^*UV1*^ mutant genomic DNA identified a region on the left arm of linkage group I that is enriched for Oak Ridge single nucleotide polymorphisms (SNPs) and contains a premature stop codon in the BAH domain of NCU07505 (BAH domain, light blue; PHD finger (split), dark blue; no annotated domains, gray). Each translucent point represents a running average of SNPs (window size = 10 SNPs, step size = 1 SNP). **f**, Serial dilution spot test silencing assay for the indicated strains. *epr-1*^*UV1*^ + *NCU07505*^*WT*^ has a wild-type copy of *NCU07505* at the *his-3* locus. All strains harbor *P*_*NCU05173*_*::hph*.

### Mapping and identification of *epr-1* as *NCU07505*

In order to map and identify the causative mutation in the *epr-1*^*UV1*^ mutant, we crossed *epr-1*^*UV1*^, which is in an Oak Ridge genetic background, to a highly polymorphic wild-type strain named “Mauriceville” (27). We then pooled the genomic DNA from Hygromycin B-resistant progeny and subjected it to whole-genome sequencing (∼15x coverage; Fig. 1d). When we scored the percentage of Oak Ridge single nucleotide polymorphisms (SNPs) across the genome (28), we found a region on linkage group (LG) I that was enriched for Oak Ridge SNPs and included an early stop mutation (Q206*, CAG->TAG) in *NCU07505* (Fig. 1e).

To verify that the early stop in *NCU07505* is the causative mutation in *epr-1*^*UV1*^, we targeted a wild-type copy of *NCU07505* to the *his-3* locus in the *epr-1*^*UV1*^ mutant background. This ectopic copy of *NCU07505* complemented the mutation, *i*.*e*., it restored drug sensitivity (Fig. 1f). In addition, we found that deletion of *NCU07505* resulted in resistance to Hygromycin B, similar to the *epr-1*^*UV1*^ strain (Fig. 1f). We subsequently isolated and characterized three additional alleles of *epr-1* generated in the mutagenesis, further supporting the notion that mutations in *NCU07505* support drug resistance (Supplementary Fig. 1).

### EPR-1 and SET-7 repress an overlapping set of H3K27-methylated genes

Although our selection was designed to isolate mutants with specific defects in Polycomb repression, we could conceivably recover mutants that globally altered transcription or led to antibiotic resistance independent of *hph* or *nat-1* upregulation. To determine if EPR-1 was specifically required for repression of H3K27-methylated genes, we performed mRNA-seq on Δ*epr-1* siblings and compared the gene expression profile to previously published wild-type and Δ*set-7* data sets (23). We found that 632 genes were upregulated and 974 genes were downregulated greater than two-fold in Δ*epr-1* strains compared to wild-type strains (*P* < 0.05) (Supplementary Fig. 2). The upregulated gene set in Δ*epr-1* was significantly enriched for H3K27-methylated genes (*X*^*2*^_(1, N = 632)_ = 40.8, *P* = 1.684 × 10^−10^)(Supplementary Fig. 2), and H3K27-methylated genes upregulated in both Δ*epr-1* and Δ*set-7* significantly overlapped (*P* = 1.436 × 10^−16^) (Fig. 2a). To verify our mRNA-seq results, we performed reverse transcription followed by quantitative polymerase chain reaction (RT-qPCR) on RNA isolated from biological triplicates of wild-type, Δ*set-7*, and Δ*epr-1* strains. Five of six examined H3K27-methylated genes found upregulated in both Δ*set-7* and Δ*epr-1* strains by mRNA-seq were confirmed by RT-qPCR (Fig. 2b and Supplementary Fig. 2). In contrast, only one out of six H3K27-methylated genes found exclusively upregulated in Δ*set-7* or Δ*epr-1* by mRNA-seq was confirmed by RT-qPCR (Supplementary Fig. 2). Thus, these data show that loss of EPR-1 derepresses a significant number of H3K27-methylated genes that are also upregulated in strains lacking H3K27 methylation.

**Fig. 2.**
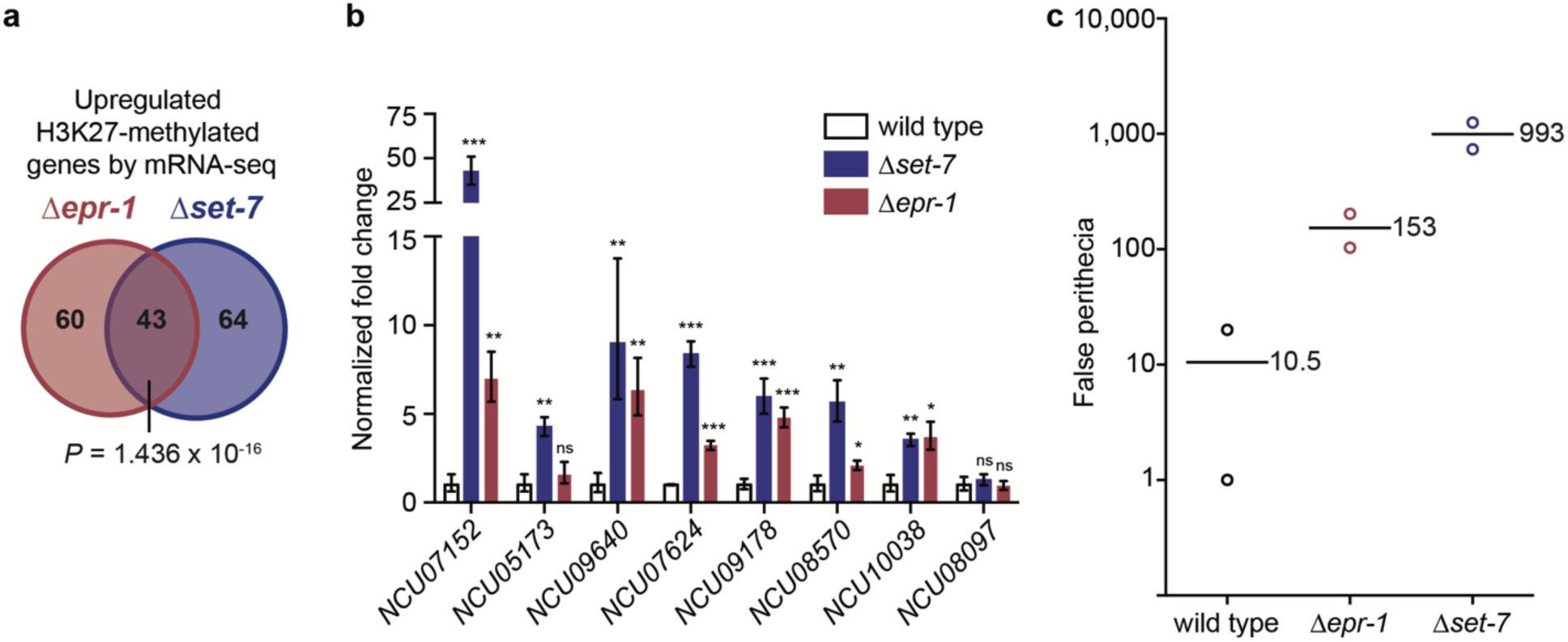
*Δepr-1* and *Δset-7* strains share defects in transcriptional silencing and sexual development. **a**, Venn diagram depicting H3K27-methylated genes that appear upregulated by mRNA-seq in both *Δepr-1* and *Δset-7* strains, only in *Δepr-1* strains, or only in *Δset-7* strains, using a significance cutoff of log_2_(mutant/wild type) > 1 and a *P* value < 0.05 using the Benjamin-Hochberg correction for multiple comparisons. Significance of genes upregulated in both *Δepr-1* and *Δset-7* strains was determined using a hypergeometric test. **b**, RT-qPCR of H3K27-methylated genes that were replaced with antibiotic resistance genes (*NCU07152, NCU05173*) and used for initial selection of mutants, and H3K27-methylated genes that appeared upregulated in both Δ*epr-1* and Δ*set-7* strains by mRNA-seq (*NCU09640, NCU07624, NCU09178, NCU08570, NCU010038, NCU08097*). Each value was normalized to expression of actin gene (*act*) and presented relative to wild type. Filled bars represent the mean from biological triplicates and error bars show standard deviation. (*** for *P* < 0.001, ** for *P* < 0.01, * for *P* < 0.05, and ns for not significant; all relative to wild type by two-tailed, unpaired t-test). **c**, Quantification of false perithecia developed in a Petri dish (85 mm diameter) after two weeks of unfertilized growth are shown for the indicated strains. Horizontal lines and numbers indicate the mean of two biological replicates (open circles).

### Δ*epr-1* and Δ*set-7* strains share a sexual development defect

Since Δ*epr-1* and *Δset-7* strains exhibit similar transcriptional profiles, we wondered if they also shared vegetative growth and sexual development phenotypes as well. To assess if *Δepr-1* strains have an altered vegetative growth rate, we measured linear growth rates of wild-type, Δ*set-7* and Δ*epr-1* strains with ‘race tubes’ (29). We confirmed that Δ*set-7* strains do not have a linear growth defect(22) and found that Δ*epr-1* strains also grow at wild-type rates (Supplementary Fig. 2). Loss of SET-7 has been implicated in promoting sexual development in mutants that are homozygous sterile (30). To determine if Δ*set-7* and/or Δ*epr-1* strains aberrantly promote sexual differentiation in the absence of a mating partner, we singly inoculated crossing plates (31) with wild-type, Δ*set-7* or Δ*epr-1* strains. After two weeks of unfertilized growth at 25 °C, we observed the development of few false perithecia with wild-type controls, whereas Δ*epr-1* and Δ*set-7* developed approximately 10- and 100-fold more false perithecia than wild type, respectively (Fig. 2c and Supplementary Fig. 2). These data suggest that EPR-1, and SET-7 to a greater extent, repress premature sexual development, which is reminiscent of fertilization-independent seed development observed in plant Polycomb mutants (32, 33).

### H3K27 methylation is essentially normal in Δ*epr-1*

As a first step to assess if the transcriptional silencing and sexual development defects shared between Δ*epr-1* and Δ*set-7* strains were due to a common global loss of H3K27 methylation, we performed a western blot on whole cell lysates to detect H3K27me3 in wild-type, Δ*set-7*, and Δ*epr-1* strains. The total levels of H3K27me3 in Δ*epr-1* strains were comparable to that in wild type (Fig. 3a and Supplementary Fig. 3). However, because only a subset of H3K27-methylated genes are derepressed in both Δ*epr-1* and Δ*set-7* strains, we wanted to know if H3K27 methylation might be specifically lost at upregulated genes in Δ*epr-1* strains. To examine this possibility, we performed H3K27me2/3 chromatin immunoprecipitation followed by sequencing (ChIP-seq) on two Δ*epr-1* siblings and compared the data to that for wild type(34). We found that the global distribution of H3K27me2/3 in Δ*epr-1* appeared to mirror that of wild type (Fig. 3b). Comparison of H3K27me2/3 levels associated with individual genes showed good agreement between the averaged wild-type and Δ*epr-1* data sets (R^2^ = 0.9105), and within replicate data for wild-type (R^2^ = 0.9272) and Δ*epr-1* (R^2^ = 0.8941) strains (Fig. 3c and Supplementary Fig. 3). We did, however, identify 29 genes with a greater than two-fold decrease and nine genes with a greater than two-fold increase in H3K27me2/3 levels in Δ*epr-1* compared to wild type. Interestingly, none of these 38 genes with altered H3K27 methylation were classified among the upregulated or downregulated gene sets in the mRNA-seq analysis of Δ*epr-1*. To validate the H3K27me2/3 ChIP-seq results, we performed H3K27me2/3 ChIP followed by quantitative Polymerase Chain Reaction (ChIP-qPCR) on wild-type, Δ*set-7*, and Δ*epr-1* strains in biological triplicate. These data confirmed wild-type levels of H3K27me2/3 at a subtelomere (Tel IL) and at the genes replaced by the antibiotic-resistance markers (*NCU07152* and *NCU05173*), and also corroborated the loss (*NCU08834*) and gain (*NCU02856*) of H3K27me2/3 observed in the ChIP-seq of Δ*epr-1* strains (Fig. 3d). Altogether, these data show that the derepression of H3K27-methylated genes in Δ*epr-1* strains is not due to concomitant loss of H3K27 methylation.

**Fig. 3.**
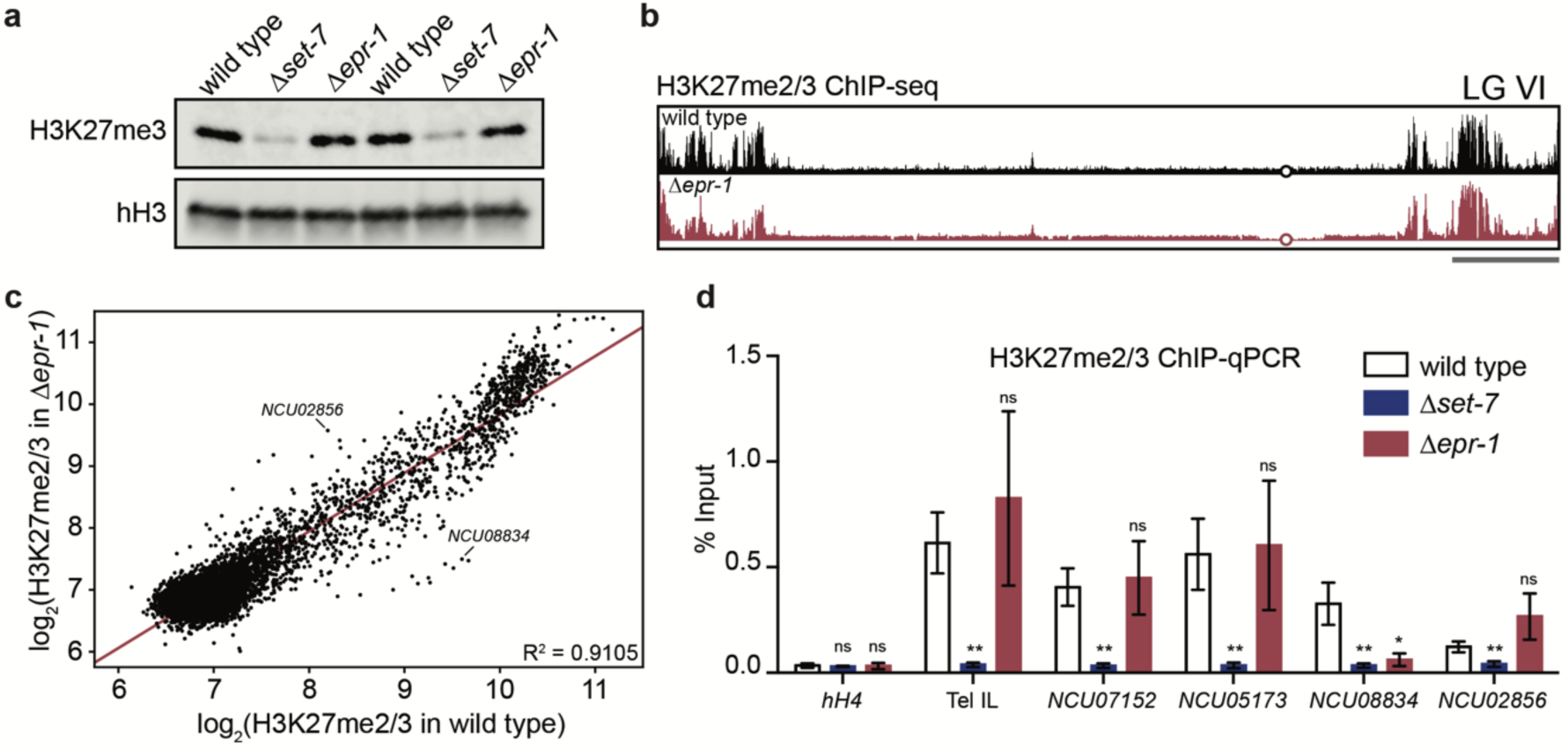
*epr-1* is not required for H3K27 methylation. **a**, Western blot showing H3K27me3 and total histone H3 (hH3) in the indicated strains. Biological replicates are shown. The same lysate was run on separate gels, and hH3 was used as a sample processing control. **b**, ChIP-seq track showing average levels of H3K27me2/3 from two biological replicates of wild-type and *Δepr-1* strains on LG VI. Open circle indicates the middle of the centromere region. Gray bar represents 500 kb. Y-axis is 0-800 RPKM for wild type and 0-1200 RPKM for *Δepr-*1. **c**, Scatter plot showing the correlation of H3K27me2/3 levels at all genes (black dots) in wild-type and *Δepr-1* strains based on biological replicates of ChIP-seq data. Line of best fit displayed in red (R^2^ = 0.9105). Representative genes that gained (*NCU02856*) or lost (*NCU08834*) H3K27me2/3 in *Δepr-1* are indicated. **d**, H3K27me2/3 ChIP-qPCR to confirm ChIP-seq data at six regions in wild-type, Δ*set-7* and Δ*epr-1* strains: *hH4* (negative control), Tel IL (unchanged H3K27me2/3 in *Δepr-1*), *NCU07152* (unchanged H3K27me2/3 in *Δepr-1*), *NCU05173* (unchanged H3K27me2/3 in *Δepr-1*), *NCU08834* (loss of H3K27me2/3 in *Δepr-1*) and *NCU02856* (gain of H3K27me2/3 in *Δepr-1*). Filled bars represent the mean of biological triplicates and error bars show standard deviation (** for *P* < 0.01, * for *P* < 0.05, and ns for not significant; all relative to wild type by two-tailed, unpaired t-test).

### EPR-1 forms telomere-associated foci dependent on EED

To localize EPR-1 *in vivo*, we used the *ccg-1* promoter to drive expression of wild-type EPR-1 fused with GFP at its N-terminus (EPR-1^WT^) (35) in a strain that had fluorescent markers for the nuclear membrane (ISH1), telomeres (TRF1), and centromeres (CenH3) (23). We found that EPR-1^WT^ was restricted to the nucleus and formed distinct foci that were typically closely associated with TRF1 foci (Fig. 4a). EPR-1^WT^ foci were significantly closer to telomeres as compared to centromeres (negative control; *P* = 0.0403) (Supplementary Fig. 4), and the number of EPR-1^WT^ and TRF1 foci per nucleus were not statistically different (*P* = 0.7422), although the majority of nuclei examined had more EPR-1^WT^ than TRF1 foci (Supplementary Fig. 4). Considering that H3K27 methylation is predominantly present near chromosome ends in *N. crassa*(22), it is not unexpected that a putative PcG protein, such as EPR-1, would co-localize with the telomere marker, TRF1.

**Fig. 4.**
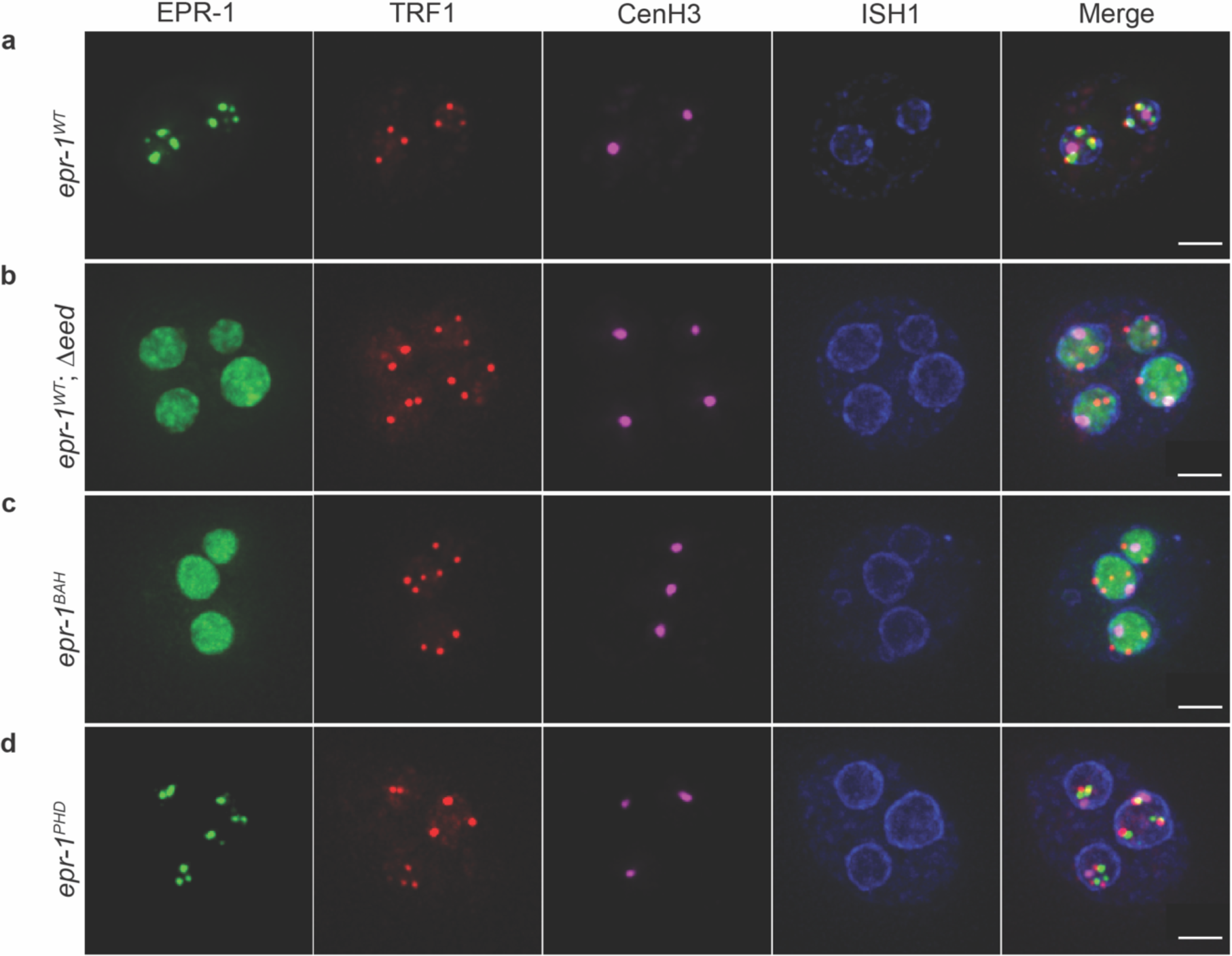
EPR-1 forms telomere-associated foci that are dependent on EED and the BAH domain of EPR-1. Maximum intensity projection images of fluorescence microscopy Z-stacks showing EPR-1 (GFP-EPR-1, green) for *epr-1*^*WT*^ (**a**), *epr-1*^*WT*^; *Δeed* (**b**), *epr-1*^*BAH*^ (**c**), and *epr-1*^*PHD*^ (**d**). Telomeres (TRF1*-*TagRFP-T, red), centromeres (CenH3-iRFP670, magenta), the nuclear membrane (ISH1*-*TagBFP2, blue), and merged images are shown for reference. Each image shows a single conidium with multiple nuclei. Overlaid white bar represents 2 µm.

To determine if the formation of EPR-1^WT^ foci was dependent on H3K27 methylation, we introduced a deletion of *eed*, a gene encoding a component of PRC2 necessary for catalytic activity(22). Strains bearing this deletion lacked distinct nuclear foci of EPR-1^WT^ and instead displayed a diffuse nuclear distribution of EPR-1^WT^ (Fig. 4b). Thus, an intact PRC2 complex or H3K27 methylation is required for proper EPR-1^WT^ subnuclear localization.

### An intact BAH domain is required for normal nuclear distribution of EPR-1

EPR-1 is predicted (36) to have a BAH domain and PHD finger, protein modules implicated in chromatin engagement (37, 38). To determine if these domains are necessary for the formation of the EPR-1^WT^ foci, we created GFP-EPR-1 constructs in which a previously identified critical tryptophan in either the BAH domain (EPR-1^BAH^) or PHD finger (EPR-1^PHD^) was replaced with an alanine (39, 40). We found that EPR-1^BAH^ displayed the same diffuse nuclear distribution as EPR-1^WT^ in a Δ*eed* background, consistent with the possibility that the BAH domain mediates interaction with H3K27-methylated chromatin (Fig. 4c). In contrast, EPR-1^PHD^ still formed nuclear foci that were equivalent to EPR-1^WT^ in number and proximity to TRF1 foci (Fig. 4d and Supplementary Fig. 4), demonstrating the nonessential nature of this conserved tryptophan residue for the normal nuclear distribution of EPR-1.

### EPR-1 localizes to H3K27 methylation genome-wide

Since proper subnuclear localization of EPR-1^WT^ appeared to require H3K27 methylation, we performed ChIP-seq to determine if EPR-1^WT^ genomic targets coincided with H3K27me2/3 throughout the genome (Fig. 5a). Results of EPR-1^WT^ ChIP-seq appeared to match the distribution of H3K27me2/3 in wild type (Fig. 5a) and we found good correlation between EPR-1^WT^ and H3K27me2/3 relative sequencing coverage over each gene (R^2^ = 0.8675) (Supplementary Fig. 5). We also examined the genomic distribution of EPR-1^PHD^ and EPR-1^BAH^ mutant alleles (Fig. 5a), which were expressed at comparable levels (Supplementary Fig. 5). The ChIP-seq coverage of EPR-1^PHD^ was still enriched at H3K27-methylated genes, albeit less so than EPR-1^WT^, while the EPR-1^BAH^ ChIP-seq did not show enrichment (Supplementary Fig. 5). To validate the ChIP-seq of GFP-EPR-1-expressing strains, we performed ChIP-qPCR for representative regions (Fig. 5b). Consistent with the ChIP-seq data, EPR-1^WT^, and EPR-1^PHD^ to a lesser degree, were enriched at examined regions bearing H3K27 methylation. In addition, the ChIP-qPCR confirmed that EPR-1^BAH^, as well as EPR-1^WT^ in a Δ*eed* background, lack such enrichment.

**Fig. 5.**
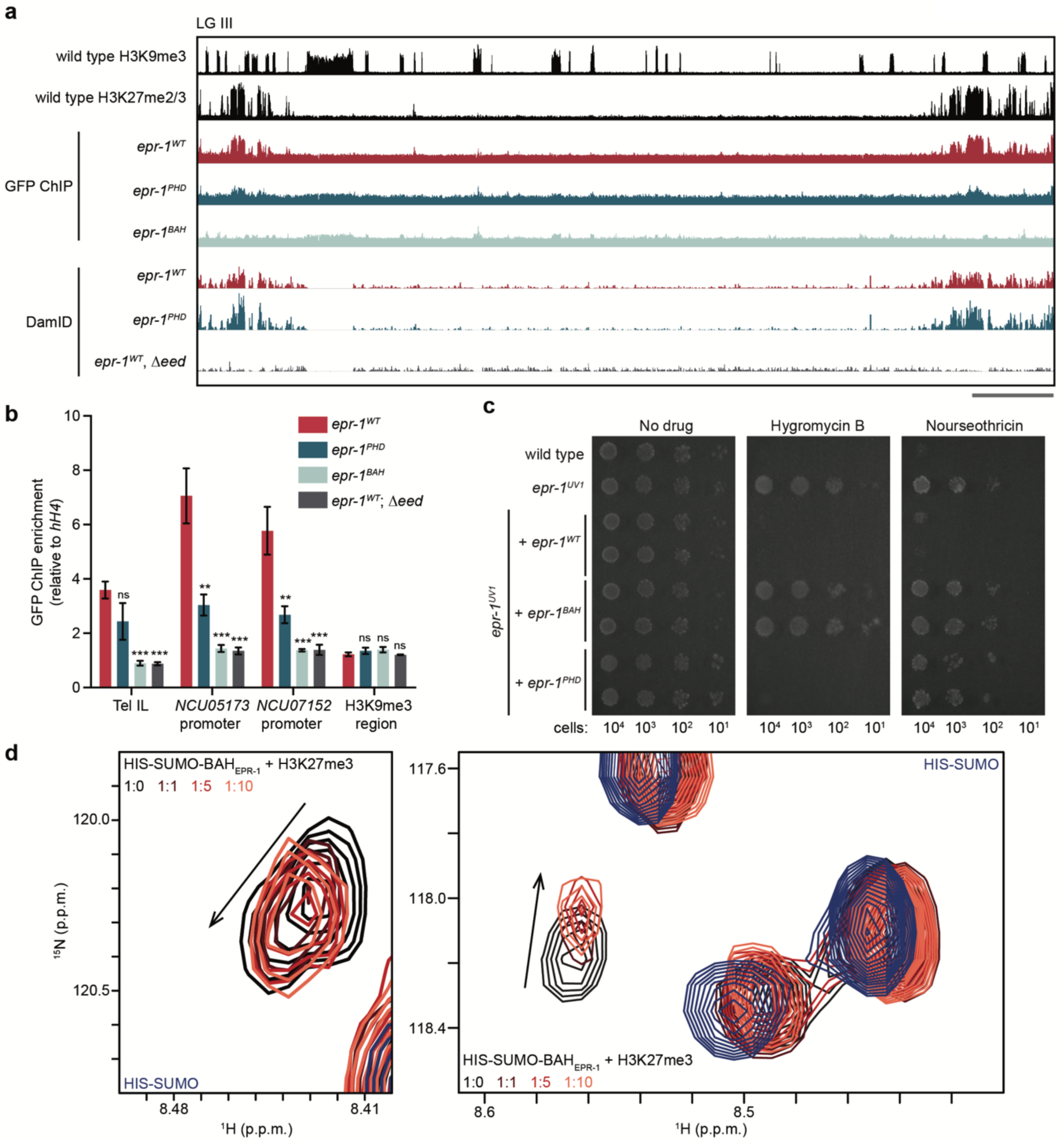
EPR-1 directly interacts with H3K27 methylated chromatin through its BAH domain. **a**, ChIP-seq and DamID-seq tracks showing average levels from two biological replicates for the indicated genotypes on LG III. Y-axis is 0-800 RPKM for H3K9me3 and H3K27me2/3 ChIP-seq, 0-500 RPKM for all GFP ChIP-seq, and 0-3500 RPKM for all DamID-seq. Gray bar represents 500 kb. **b**, GFP ChIP-qPCR to validate GFP-EPR-1 ChIP-seq data at four genomic regions: Tel IL (H3K27-methylated), *NCU05173* promoter (H3K27-methylated), *NCU07152* promoter (H3K27-methylated), and H3K9me3 region (negative control, LG VI centromere). All data are normalized to a negative, euchromatic control, *hH4*. Filled bars represent the mean and error bars show standard deviation from three biological replicates (*** for *P* < 0.001, ** for *P* < 0.01, and ns for not significant; all relative to wild type by two-tailed, unpaired t-test). **c**, Serial dilution spot test silencing assay for the indicated strains. All strains harbor *P*_*NCU05173*_*::hph* and *P*_*NCU07152*_*::nat-1*. **d**, Two regions of an overlay of ^1^H-^15^N HSQC spectra of ^15^N-labelled HIS-SUMO-BAH_EPR-1_ fusion in the presence of increasing concentrations of H3K27me3 peptide. Spectra are color coded as indicated. ^15^N-labelled HIS-SUMO is included for reference.

As an orthogonal approach to ChIP, we determined the chromatin targets of EPR-1 by fusing an *E. coli* DNA adenine methyltransferase (Dam) (41) to the C-terminus of endogenous EPR-1 and assayed adenine-methylated DNA fragments by sequencing (DamID-seq) (42) (Fig. 5a). Using DamID-seq of EPR-1^WT^-Dam and methyl-sensitive restriction enzyme Southern blots, we found that EPR-1^WT^-Dam localizes to H3K27-methylated genes and this is dependent upon EED (Fig. 5a and Supplementary Fig. 5). Mutation of the PHD finger in the EPR-1-Dam fusion did not abolish its targeting to H3K27-methylated chromatin (Fig. 5a). These results were consistent with our ChIP-seq findings. We conclude that EPR-1 localizes to H3K27-methylated regions of the genome and that proper recruitment of EPR-1 to chromatin requires both an intact BAH domain and the integral PRC2 component, EED.

### Both the BAH domain and PHD finger of EPR-1 are necessary for gene repression

Our localization studies of EPR-1^PHD^ and EPR-1^BAH^, while suggestive, did not directly test the role of the PHD finger and BAH domain of EPR-1 in H3K27 methylation-mediated silencing. We therefore utilized our antibiotic-resistance reporters, used in the initial selection, to test more directly their possible involvement in gene repression. We targeted ectopic copies of *epr-1*^*WT*^, *epr-1*^*BAH*^ or *epr-1*^*PHD*^ to the *his-3* locus in an *epr-1*^*UV1*^ strain bearing the antibiotic-resistance genes and scored drug resistance. Whereas *epr-1*^*WT*^ restored sensitivity to Hygromycin B and Nourseothricin, *epr-1*^*BAH*^ remained resistant to both drugs (Fig. 5c). In contrast, introduction of *epr-1*^*PHD*^ apparently re-silenced the *hph*, but not the *nat-1*, antibiotic-resistance gene (Fig. 5c). This suggests that while the PHD finger of EPR-1 is not essential for recruitment to H3K27 methylated chromatin, it is not entirely dispensable for gene silencing.

### BAH domain of EPR-1 binds to H3K27me3 *in vitro*

To test, directly, whether the BAH domain of EPR-1 recognizes H3K27 methylation, we cloned, expressed, and purified a HIS-SUMO-BAH_EPR-1_ fusion protein for use in nuclear magnetic resonance (NMR) spectroscopy experiments. We found that the BAH_EPR-1_ domain alone was unstable once cleaved from the HIS-SUMO tag and therefore we used the HIS-SUMO-tagged BAH_EPR-1_ in all subsequent experiments. An initial ^1^H–^15^N-heteronuclear single quantum coherence (^1^H–^15^N HSQC) spectrum revealed that the fusion protein was well folded (Supplementary Fig. 5). An overlay with an ^1^H–^15^N HSQC spectrum of ^15^N labelled HIS-SUMO tag alone was used to determine which peaks belonged to BAH_EPR-1_ (Supplementary Fig. 5). Addition of increasing concentrations of an H3K27me3 peptide to the HIS-SUMO-BAH_EPR-1_ fusion protein led to significant chemical shift perturbations (CSPs), indicating binding (Fig. 5d,e). Importantly, the perturbed resonances belonged exclusively to BAH_EPR-1_. We conclude that the BAH domain of EPR-1 associates with an H3K27me3 peptide. Due to protein stability problems, we were unable to calculate an accurate dissociation constant (K_d_); however, the CSPs appeared consistent with a high micromolar K_d_.

### EPR-1 is a homolog of plant EBS/SHL and widely distributed across eukaryotes

To determine if EPR-1 orthologs exist outside of *N. crassa*, we performed sequence similarity searches to identify homologs, followed by phylogenetic and domain architectural analysis of those to identify genuine orthologs. Consequently, we were able to identify orthologs in various fungal species as well as a wide range of other eukaryotes (Fig. 6a). Notably, we determined the *Arabidopsis thaliana* paralogs EBS and SHL as orthologs of EPR-1 in our analyses, which have been erroneously reported as plant-unique proteins (21, 25, 26). Similar to EPR-1, the plant paralogs, EBS and SHL, bind H3K27 methylation and have been implicated in gene repression (17, 18, 21). As one approach to investigate if other EPR-1 orthologs may have roles independent of H3K27 methylation, we checked if any species has an EPR-1 ortholog but lacks a SET-7 (H3K27 methyltransferase) ortholog. With the exception of Chytridiomycota and Fonticula lineages, in which the presence of SET-7 homologs was deemed ambiguous due to lack of a definitive pre-SET domain, all examined species with EPR-1 homologs had clear SET-7 homologs (Fig. 6a). This result is consistent with EPR-1 orthologs mediating H3K27 methylation-based repression in a wide variety of species.

**Fig. 6.**
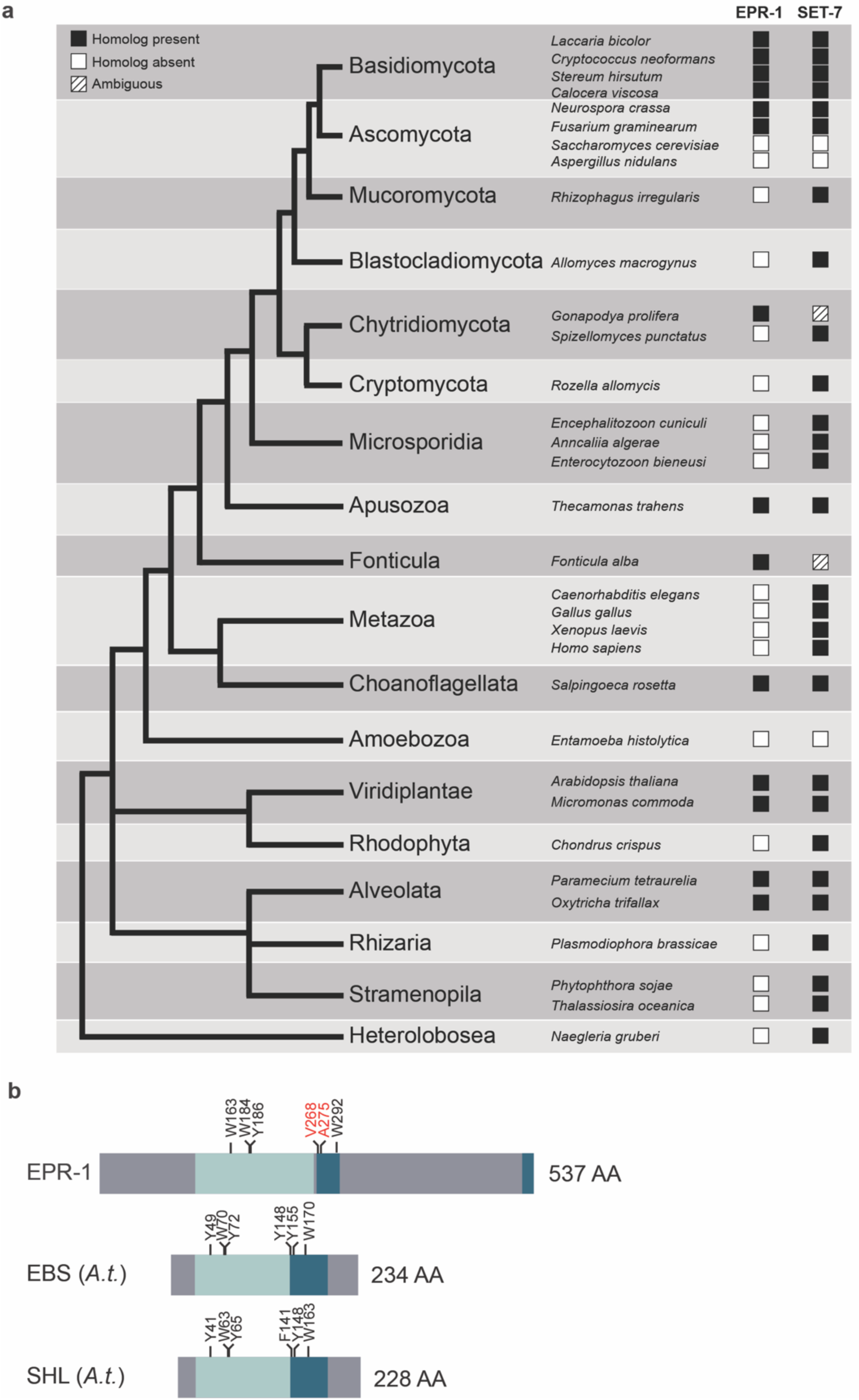
EPR-1 homologs are present in species beyond fungi. **a**, The presence and absence of EPR-1 and SET-7 protein homologs across major species divisions is depicted in a representative tree of eukaryotes. The leaves of the tree are labelled with the names of the divisions. Representative species are featured and their associated squares in the EPR-1 and SET-7 columns indicate the presence or absence of homologs, as indicated. **b**, Protein domain structure of EPR-1, as well as EBS and SHL from *Arabidopsis thaliana* (*A*.*t*.). BAH domains are indicated by light blue, the PHD fingers by dark blue, and regions with no known domains are gray. Aromatic amino acid residues involved in methylated histone recognition in the BAH domain and PHD finger are indicated above the protein structure diagram (black text). Red text above the PHD finger in EPR-1 highlights the absence of aromatic residues at these amino acid positions.

In contrast to EPR-1, *A. thaliana* EBS is not entirely restricted to H3K27-methylated genes. Indeed, the majority of EBS-bound genes are devoid of H3K27me3 and instead are associated with an ‘active’ chromatin mark, H3K4me3 (43), via the PHD finger of EBS (17). To understand this discrepancy, we examined the underlying protein sequence of the PHD finger of EPR-1. Whereas aromatic residues implicated in methylated histone recognition in the BAH domain (44) are present in both EPR-1 and the plant paralogs, comparable residues in the PHD finger (18, 45, 46) are present in EBS and SHL, but lacking in EPR-1 (Fig. 6b, highlighted in red). Furthermore, a single amino acid substitution replacing an aromatic tyrosine residue with an alanine residue in the PHD finger of a plant SHL was sufficient to diminish *in vitro* H3K4me3-binding greater than two-fold (18). Intriguingly, this residue corresponds to the naturally occurring alanine 275 in EPR-1 (Fig. 6b). This suggests that while many species have EPR-1 homologs, they may not necessarily be bivalent histone readers like plant EBS and SHL.

## Discussion

Deciphering the basic mechanism(s) of Polycomb repression has been difficult, in part, due to the diversity (47), redundancy (48), as well as interdependence (15) of protein players involved. For this reason, the model organism *N. crassa*, which employs H3K27 methylation catalyzed by PRC2 for gene repression (22), yet conspicuously lacks PRC1 components (7), represents an ideal organism to uncover fundamental aspects of Polycomb silencing. Here we have identified, to the best of our knowledge, the first known reader and effector of H3K27 methylation in fungi, EPR-1. This provides insight into how Polycomb silencing can function in the absence of PRC1.

Our ChIP and DamID results demonstrated that EPR-1 co-localizes with H3K27 methylation genome-wide and cytological examination revealed that GFP-EPR-1 forms approximately 3-5 foci per nucleus. This implies that domains of H3K27 methylation within and between the seven *N. crassa* chromosomes generally self-associate. This is consistent with the observed intra- and inter-chromosomal contacts among H3K27-methylated regions of the genome in Hi-C experiments (49). Similar, and perhaps equivalent, higher-order chromatin structures, referred to as Polycomb bodies (24), are known to be mediated by PRC1 components in both plant and animal cells (50-53). Our group has previously shown that SET-7, the catalytic component of PRC2, is required for normal 3D genome organization (23). It would be interesting to learn if EPR-1, the only known effector of H3K27 methylation in *N. crassa*, is also essential for this wild-type chromatin organization.

Despite the loss of transcriptional silencing observed in *epr-1* mutants, they do not exhibit appreciably altered H3K27 methylation – this is striking for a few reasons. First, it suggests that H3K27 methylation-mediated silencing is a unidirectional pathway in *N. crassa*, in which EPR-1 acts downstream of PRC2. This is in contrast to findings in plants and animals, in which PRC1 components can affect the recruitment or activity of PRC2 (21, 54, 55), a fact that has hampered the elucidation of the direct role of PRC components in gene repression. Second, since Δ*epr-1* strains de-repress H3K27-methylated genes without loss of H3K27 methylation, it suggests that active transcription does not necessarily preclude PRC2 activity. This was surprising since transcriptional shut-off is thought to precede PRC2 activity during normal animal development (56) and because artificial gene repression can be sufficient to recruit PRC2 (57, 58). Finally, the presence of H3K27 methylation on de-repressed genes in Δ*epr-1* strains demonstrates that H3K27 methylation *per se* is not sufficient for effective gene repression, consistent with previous reports (59, 60). It is noteworthy, however, that while Δ*set-7* and Δ*epr-1* strains share transcriptional and sexual defects, the phenotype of Δ*set-7* strains is generally more pronounced. This suggests that PRC2 or H3K27 methylation may have additional roles in gene repression that go beyond recruitment of EPR-1 to chromatin.

Our investigation into the phylogenetic distribution of EPR-1 homologs indicates that an ancestral EPR-1 emerged prior to the divergence of plants, animals, and fungi. This ancestral EPR-1 may have been an integral component of an early eukaryotic Polycomb silencing system. While animals are a notable exception, regarding the absence of EPR-1 homologs with a BAH-PHD structure, a human BAH domain-containing protein, BAHD1, has been reported to ‘read’ H3K27me3 (39) and promote gene silencing (61), although apparently not interact with known PRC1 components (62). It is therefore conceivable that BAHD1 homologs present in animal lineages are actually divergent orthologs of EPR-1 that lack a PHD finger. Regardless of their ancestry, both human BAHD1 and *N. crassa* EPR-1 represent novel forms of H3K27 methylation-mediated repression that do not rely on PRC1 components.

## Materials and Methods

### Strains, media and growth conditions

All *N. crassa* strains used in this study are listed in Supplementary Table 1. Liquid cultures were grown with shaking at 32 °C in Vogel’s minimal medium (VMM) with 1.5% sucrose (63). Crosses were performed at 25 °C on modified Vogel’s with 0.1% sucrose (31). Spot tests were performed at 32 °C on VMM with 0.8% sorbose, 0.2% fructose, and 0.2% glucose (FGS). When appropriate, plates included 200 µg/mL Hygromycin B Gold (InvivoGen) or 133 µg/mL Nourseothricin (Gold Biotechnology). Linear growth rates were determined as previously described except 25 mL serological pipettes were used in place of glass tubes (29). Genomic DNA was isolated as previously described (64). Nutritional supplements required for auxotrophic strains were included in all growth media when necessary.

### Selection for mutants defective in H3K27 methylation-mediated silencing

Ten thousand conidia of strain N6279 (created with primers in Supplementary Table 2) were plated on VMM supplemented with FGS and 500 µg/mL histidine and subjected to 0, 3, 6, or 9 seconds of ultraviolet light (Spectrolinker XL-1500 UV Crosslinker, Spectronics Corporation) in a dark room and plates were wrapped in aluminum foil to prevent photoreactivation (65). Plates were incubated at 32 °C for 16 hours before being overlaid with 1% top agar containing VMM, FGS, 500 µg/mL histidine, Hygromycin B and Nourseothricin. Drug-resistant colonies were picked after an additional 48-72 hours at 32 °C. Initial mutant strains were crossed to a *Sad-1* mutant (66) strain (N3756) and resultant progeny were germinated on Hygromycin B- and Nourseothricin-containing medium to obtain homokaryotic mutants.

### Whole genome sequencing, mapping and identification of *epr-1* alleles

Homokaryotic mutants resistant to Hygromycin B and Nourseothricin were crossed to the genetically polymorphic Mauriceville strain (FGSC 2225) (27) and resultant progeny were germinated on medium containing Hygromycin B and/or Nourseothricin to select for strains bearing the causative mutation. Genomic DNA from approximately 15-20 progeny per mutant were pooled and sequencing libraries were prepared with a Nextera kit (Illumina, FC-121-1030). All whole genome sequencing data is available on NCBI SRA (accession #PRJNA526508). To map the approximate location of a particular causative mutation, the fraction of Oak Ridge (versus Mauriceville) SNPs across the *N. crassa* genome was determined as previously described (28) and visualized as a moving average (window size = 10 SNPs, step size = 1 SNP) with Matplotlib (67). We utilized FreeBayes and VCFtools to identify genetic variants present in our pooled mutant genomic DNA but absent in the original mutagenized strain (N6279) and the Mauriceville strain (68, 69). Only genetic variants of high probability that were consistent with the mapping data were considered further.

### Western blotting

*N. crassa* tissue from a 16 hour liquid culture of germinated conidia was collected by filtration, washed with 1x phosphate-buffered saline (PBS; 137 mM NaCl, 10 mM phosphate, 2.7 mM KCl, pH 7.5), and suspended in 500 μL of ice-cold lysis buffer (50 mM HEPES [pH 7.5], 150 mM NaCl, 10% glycerol, 0.02-0.2% NP-40, 1 mM EDTA) supplemented with 1x Halt™ Protease Inhibitor Cocktail (Thermo Scientific). Tissue was sonicated (Branson Sonifier-450) for three sets of 10 pulses (Output = 2, Duty cycle = 80), keeping the sample on ice between sets. Insoluble material was pelleted by centrifugation at 14,000 RPM at 4 °C for 10 minutes and the supernatant used as the western sample. Anti-H3K27me3 (Cell Signaling Technology, 9733) and anti-hH3 (Abcam, ab1791) primary antibodies were used with IRDye 680RD goat anti-rabbit secondary (LI-COR, 926-68071). Anti-GFP (Thermo Fisher, A10262) primary antibody was used with goat anti-chicken HRP conjugated secondary antibody (Abcam, 6877). Images were acquired with an Odyssey Fc Imaging System (LI-COR) and analyzed with Image Studio software (LI-COR).

### Chromatin immunoprecipitation

Liquid cultures were grown for approximately 18 hours with shaking at 32 °C. Tissue samples for H3K27me2/3 ChIP were cross-linked with 0.5% formaldehyde for 10 minutes and GFP ChIP samples were cross-linked with 1% formaldehyde for 10 minutes in 1x PBS. Cross-linking was quenched with 125mM glycine, tissue was washed with 1x PBS and collected. Cells were lysed in ChIP lysis buffer (50 mM HEPES [pH 7.5], 90 mM NaCl, 1 mM EDTA, 1% Triton X-100, 0.1% Deoxycholate) supplemented with 1x Halt protease inhibitor cocktail (Thermo Scientific) using a Branson Sonifier 450. Chromatin was sheared using a Bioruptor (Diagenode) and 2 µl of appropriate antibody (H3K27me2/3, Active Motif 39536, or GFP, MBL 598) was added and incubated with rotation at 4 °C overnight. Protein A/G agarose (40 µL; Santa Cruz Biotechnologies) was added to H3K27me2/3 ChIP samples and Protein A agarose (40 µL; Sigma) was added to GFP ChIP samples and incubated for 3 hours, rotating at 4 °C. Beads were washed twice with ChIP lysis buffer supplemented with 140 mM NaCl, once with ChIP lysis buffer with 0.5 M NaCl, once with LiCl wash buffer (10 mM Tris-HCl [pH 8.0], 250 mM LiCl, 0.5% NP-40, 0.5% Deoxycholate, 1 mM EDTA), and once with TE (10 mM Tris-HCl, 1 mM EDTA), all rotating at 4 °C for 10 minutes each. DNA was eluted from beads by incubation in TES (10 mM Tris-HCl, 1 mM EDTA, 1% sodium dodecyl sulfate) at 65 °C. Crosslinking was reversed by incubation at 65 °C for 16 hours and then samples were treated with proteinase K for 2 hours at 50 °C. DNA was purified using Minelute columns (Qiagen) and subsequently used for qPCR with the PerfCTa SYBR Green FastMix (QuantBio, 95071-012) on a Step One Plus Real Time PCR System (Life Technologies) using primer pairs in Supplementary Table 3, or prepared for sequencing using the NEBNext DNA Library Prep Master Mix Set for Illumina (New England BioLabs). ChIP-seq data are available on the NCBI GEO database (GSE128317).

### ChIP-seq mapping and analysis

The suite of tools available on the open-source platform Galaxy (70) was used to map ChIP-sequencing reads(71) against the corrected *N. crassa* OR74A (NC12) genome(49) and to create bigWig coverage files normalized to reads per kilobase per million mapped reads (RPKM) (72). ChIP-seq tracks were visualized with the Integrative Genomics Viewer (73). MAnorm was used to compare H3K27me2/3 ChIP-seq coverage on all genes (designated as ‘peaks’) between samples and data were visualized with Matplotlib (67). Genes were scored by their normalized H3K27me2/3 ChIP-seq coverage in wild type and the top-ranking genes (873) were designated as H3K27 methylated.

### RNA isolation, mRNA-seq library prep, and RT-qPCR

RNA was extracted from germinated conidia grown for 16-18 hours with a 1:1:1 glass beads, NETS (300mM NaCl, 1mM EDTA, 10mM Tris-HCl, 0.2% SDS), acid phenol:chloroform mixture (5:1; [pH 4.5]) using a bead beater and ethanol precipitated. RNA was treated with DNAse I (Amplification grade; Thermo Fisher Scientific). DNAse I-treated RNA was used for RNA-seq library preparation as previously described(23) or cDNA was synthesized using the SuperScript III First Strand-Synthesis System (Thermo Fisher Scientific) with poly-dT primers. cDNA was used for qPCR using the PerfCTa SYBR Green FastMix (QuantBio) on a Step One Plus Real Time PCR System (Life Technologies) using primer pairs in Supplementary Table 4. mRNA-seq data are available on the NCBI GEO database (GSE128317).

### mRNA-seq mapping and analysis

Tools available on Galaxy (70) were used to map mRNA-sequencing reads (intron size < 1kb)(74) against the corrected *N. crassa* OR74A (NC12) genome (49), to count the number of reads per gene(74) and to identify differentially expressed genes (75). *P* values reported are adjusted for false discovery rates (75). The Chi-squared test was used to determine if upregulated genes in Δ*epr-1* strains were enriched for genes marked with H3K27 methylation. Significance of gene set intersections were calculated with a hypergeometric distribution (http://nemates.org/MA/progs/overlap_stats_prog.html).

### DamID Southern hybridizations and sequencing

Southern hybridizations were carried out as previously described (76), except probes were made with PCR products amplified from wild-type *N. crassa* genomic DNA (*NCU05173*, Tel VIIL) or plasmid pBM61 (*his-3*) using primer pairs from Supplementary Table 5. Preparation of N6-methyladenine-containing DNA for sequencing was performed using a previously reported procedure (42) using primers in Supplementary Table 6 with the following modifications: 5 μg of genomic DNA from *N. crassa* strains expressing a Dam fusion was digested with 1 μL of DpnI (NEB, 20 units/μL); ligation to primer 5050 was carried out overnight at 16 °C; amplification reactions of ligated DNA with primer 5051 were performed in triplicate with 5 μL dNTPs, and an additional PCR cycle was added to the 2^nd^ and 3^rd^ phase of the PCR protocol (4 and 18 cycles respectively); 3 μg of pooled, amplified DNA was sheared using a Bioruptor (Diagenode) twice on high for 10 minutes (30 seconds on/off) at 4 °C; biotinylated DNA was purified using 250 μL slurry of streptavidin-agarose beads (Sigma). DNA was cleaved from the beads with DpnII and libraries were prepared for sequencing using the NEBNext DNA Library Prep Master Mix Set for Illumina (New England BioLabs).

### False perithecia assay and image analysis

To assay of the development of false perithecia, strains were grown on modified Vogel’s media(31) as described above without a fertilizing strain. Images of plates were acquired after two weeks of growth at 25 °C and false perithecia were detected and quantified using the Laplacian of Gaussian blob detection algorithm from scikit-image (77).

### Microscopy image acquisition and analysis

Live conidia were suspended in water and placed on a poly-L-lysine (Sigma) coated coverslip (No. 1.5; VWR) and mounted on a glass slide. Single plane images for distance measurements were captured with the ELYRA S.1 system (Zeiss) mounted on an AXIO Observer Z1 inverted microscope stand (Zeiss) equipped with a 63x, 1.4 NA Plan-Apochromat oil-immersion lens (Zeiss) and analyzed using Imaris (version 9.2.1). Images for volume renderings and max projections were collected with the DeltaVision Ultra microscope system (GE) equipped with a 100x, 1.4 NA UPlanSApo objective (Olympus). Three-dimensional Z-stack wide-field fluorescent (eGFP, TagRFP, iRFP, and TagBFP) images were captured with an sCMOS camera controlled with Acquire Ultra software. Images were processed using 10 cycles of enhanced ratio deconvolution and max projections were made using softWoRx (GE, version 7.1.0). Imaris (version 9.2.1) was used to make volume renderings and TFR1 and EPR1 foci were counted by hand.

### Cloning, expression and purification of BAH domain from EPR-1

The BAH domain of EPR-1 (amino acids 134-264) was PCR-amplified from a plasmid containing *epr-1* cDNA with primers 6681 and 6682 (Supplementary Table 7), and cloned into pE-SUMO (LifeSensors), resulting in plasmid 3410 (Supplementary Table 8). A HIS-SUMO-only control construct was generated by adding a stop codon immediately before the BAH_EPR-1_ domain in plasmid 3410. The HIS-SUMO-BAH_EPR-1_ fusion and HIS-SUMO tag constructs were expressed in *Escherichia coli* BL21 DE3 cells (New England BioLabs). Both HIS-SUMO-BAH_EPR-1_ and HIS-SUMO cultures were initially grown in 4 liters of LB media at 37 °C at 200 RPM until cultures reached an optical density of ∼ 1.0 at 600 nm and then cultures were pelleted. Cells were then re-suspended separately in 1 liter of M9 medium supplemented with ^15^N-NH_4_Cl. HIS-SUMO-BAH_EPR-1_ cultures were induced with 0.4 mM IPTG and grown for 16-18 hours at 18 °C. HIS-SUMO cultures were induced with 1 mM IPTG and grown for 16-18 hours at 25 °C. Both cultures were collected separately by centrifugation at 6,000 RPM for 20 minutes, frozen in liquid N_2_, and stored at −80 °C. The same protocol outlined below was used for the purification of both HIS-SUMO-BAH_EPR-1_ and HIS-SUMO alone.

Cells were re-suspended in lysis buffer (20 mM Tris-HCl [pH 7.5], 500 mM NaCl, 5 mM imidazole) with DNase I, EDTA-free protease inhibitors (Roche mini-tablets) and lysozyme. The cell suspension was sonicated over a period of 1 minute, alternating 1 second on and 2 seconds off. The lysate was cleared by centrifugation at 15,000 RPM for one hour at 4 °C, and the soluble fraction was loaded onto a column packed with Ni(II)-nitrilotriacetic acid agarose (Qiagen) pre-equilibrated in lysis buffer. The column was washed with 100 mL of elution buffer containing 5 mM imidazole, and the protein was eluted with buffer containing 100 mM - 200 mM imidazole. Elution fractions were then analyzed using SDS-PAGE, were pooled and concentrated using 10,000 MWCO membrane (Millipore). The protein was further purified using fast protein liquid chromatography (FPLC). Concentrated protein was loaded onto a pre-equilibrated Superdex 75 (GE Healthcare) column containing 20 mM Tris-HCl pH 7.5, 150 mM NaCl, 2 mM DTT. Size exclusion fractions were analyzed using SDS-PAGE, concentrated, and stored at −80 °C.

### NMR spectroscopy

H3K27me3 (amino acids 23–34) peptide was obtained from Anaspec. For NMR studies, peptides were re-suspended in H_2_O to a final concentration of 20 mM, and pH was adjusted to 7.0. Titration experiments of peptide into HIS-SUMO-BAH_EPR-1_ were carried out by collecting ^1^H-^15^N HSQC spectra on ^15^N labeled protein at 0.09 mM in the presence of increasing peptide concentrations. Titration points were taken at protein:peptide molar ratios of 1:0, 1:1, 1:5 and 1:10. In addition, an ^1^H-^15^N HSQC was collected on ^15^N labeled 0.1 mM HIS-SUMO tag alone. All NMR data were collected on a Bruker Avance II 800 MHz NMR spectrometer equipped with a cryoprobe. Data were processed using NMRPipe, and further analyzed using CcpNmr.

### Bioinformatic identification of EPR-1 and SET-7 homologs

PSI-BLAST (78) was used to initiate sequence similarity searches with representative sequences of EPR-1 (accession number XP_965052.2) and SET-7 (accession number XP_965043.2, residues 577-833) from *N. crassa* against the National Center for Biotechnology Information (NCBI) non-redundant (NR) database and locally maintained databases of proteins from representative species from different branches of the tree of life. HHpred (79) was used to perform profile-profile comparisons against the Protein Data Bank (PDB), Pfam and locally maintained protein sequence profiles. Sequences were clustered using BLASTCLUST (ftp://ftp.ncbi.nih.gov/blast/documents/blastclust.html). Multiple sequence alignments were generated using Kalign (80) and then adjusted manually. FastTree (81) was used to assess phylogenetic relationships among the proteins retrieved after sequence similarity searches. Customized PERL scripts were used for the analysis of domain architectures and other contextual information about protein sequences. Stringent criteria of domain-architectural concordance were used to retrieve only orthologs of all proteins under study along with grouping in phylogenetic tree analysis. In the case of SET-7, only those protein sequences that have a complete pre-SET domain followed by a SET domain were considered for analysis.

### Replacement of *NCU05173* and *NCU07152* ORFs with *hph* and *nat-1*

To delete *NCU07152*, the 5’ and 3’ regions flanking the open reading frame (ORF) were amplified from wild-type genomic DNA with primers 6385-6388 (Supplementary Table 2). The *nat-1* gene was amplified from plasmid 3237 with primers 6269 and 6270. The resulting three pieces of DNA were stitched by overlap extension using primers 6385 and 6388 and the final product was transformed into N4840. Primary transformants were selected on Nourseothricin-containing medium and crossed to a wild-type strain (N3753) to remove *Δset-7* and *Δmus-52* from the genetic background, resulting in strain N5808. To delete *NCU05173*, the 5’ and 3’ regions flanking the ORF were amplified from wild-type genomic DNA with primers 6605-6608. The *hph* gene was amplified from plasmid 2283. The 5’ flank was stitched to the 5’ portion of *hph* by overlap extension using primers 6605 and 2955, and the 3’ flank was stitched to the 3’ portion of *hph* using primers 6608 and 2954. The resulting two pieces of ‘split marker’ DNA were transformed into strain N4840. Primary transformants were selected on Hygromycin B-containing medium and crossed to N5808 to generate a homokaryotic strain with both marker genes (N6233). N6233 was crossed to N623 to introduce *his-3*, resulting in the strain used for the mutant hunt (N6279).

### Creation and targeting of *his-3*^+^::*pCCG*::N-GFP::EPR-1 plasmids

The ORF and 3’ UTR of *epr-1* were PCR-amplified from wild-type genomic DNA with primers 6416 and 6417 (Supplementary Table 9) and cloned into plasmid 2406 (35) using PacI and XbaI restriction sites. For the BAH point mutant, *epr-1*^*W184A*^, two PCR products amplified from wild-type genomic DNA with primers pairs 6416 and 6368, and 6367 and 6417 were PCR-stitched together with primers 6416 and 6417, and similarly cloned into plasmid 2406. For the PHD point mutant, *epr-1*^*W292A*^, two PCR products amplified from wild-type genomic DNA with primer pairs 6416 and 6400, and 6399 and 6417 were PCR-stitched together with primers 6416 and 6417, and cloned into plasmid 2406. Plasmids were linearized with NdeI and targeted to *his-3* in either N7451 (for complementation) or N7567 (for microscopy), as previously described (82). Primary transformants were then crossed to N7549 or N7552, respectively.

### Replacement of *epr-1* with *trpC::nat-1*

The 5’ and 3’ flanks of *epr-1* were PCR-amplified from wild-type genomic DNA with primer pairs 6401 and 6402, and 6350 and 6351, respectively (Supplementary Table 10). The 5’ and 3’ flanks were separately PCR-stitched with plasmid 3237 (source of *nat-1*) using primer pairs 6401 and 4883, and 4882 and 6351, respectively. These two ‘split-marker’ PCR products were transformed into strain N7537 and *epr-1* replacements were selected on Nourseothricin-containing medium. Primary transformants were crossed to N3752 (to generate N7567) and N6234 (to generate N7576).

### Endogenous C-terminal-tagging of EPR-1 with 10xGly::Dam

The regions immediately upstream (5’) and downstream (3’) of *epr-1*’s stop codon were PCR-amplified from wild-type genomic DNA with primer pairs 6348 and 6349, and 6350 and 6351, respectively (Supplementary Table 11). The 5’ and 3’ regions were separately PCR-stitched with plasmid 3131 (source of 10xGly::Dam::*trpC*::*nat-1*) using primer pairs 6348 and 4883, and 4882 and 6351, respectively. These two ‘split-marker’ PCR products were transformed into N2718 and ‘knock-ins’ were selected on Nourseothricin-containing medium. Primary transformants were crossed to N3752 to obtain homokaryons. To make *epr-1*^*W292A*^ fusions with Dam, the same approach was taken except the 5’ region was a PCR product stitched from two PCR products amplified from wild-type genomic DNA with primer pairs 6346 and 6400, and 6399 and 6349.

## Acknowledgments

We wish to thank C. Petell and B. Strahl for preliminary work on binding specificity of EPR-1; S. Honda for providing some *N. crassa* strains utilized in fluorescence microscopy experiments; J. Lyle, A. Leiferman and N. Meyers for help in genetic mapping of UV-generated mutants; A. Harvey for help with preliminary microscopy experiments; and A. Zemper for the chicken anti-GFP and goat anti-chicken-HRP antibodies. This work was funded by the National Institute of General Medical Sciences (GM127142 and GM093061 to E.U.S.; GM128705 to C.A.M.), the National Science Foundation (CAREER-1452411 to C.A.M.), and the American Heart Association (14POST20450071 to E.T.W.). K.J.M. was supported by the National Institutes of Health (HD007348), and G.K. and L.A. were supported by the Intramural Research Program of the National Library of Medicine, USA.

## Author contributions

E.U.S. conceived of the forward genetic selection. E.T.W. and K.J.M. performed the majority of the experiments and data analysis. T.O. performed mRNA-seq, J.M.S. performed some microscopy experiments, S.M.D. performed NMR experiments in the laboratory of C.A.M., and G.K. did phylogenetic analyses in the laboratory of L.A. K.J.M. and E.T.W. wrote the first draft of the manuscript with input and editing by E.U.S. All authors reviewed and approved the final manuscript.

## Data availability

All ChIP-seq, DamID-seq, and mRNA-seq data have been submitted to the NCBI GEO database (GSE128317). All whole-genome sequencing data have been submitted to NCBI SRA (accession #PRJNA526508). All other datasets generated during and/or analyzed during the current study are available from the corresponding author on reasonable request. All figures have raw data associated with them.

---

**Supplementary Fig. 1.**
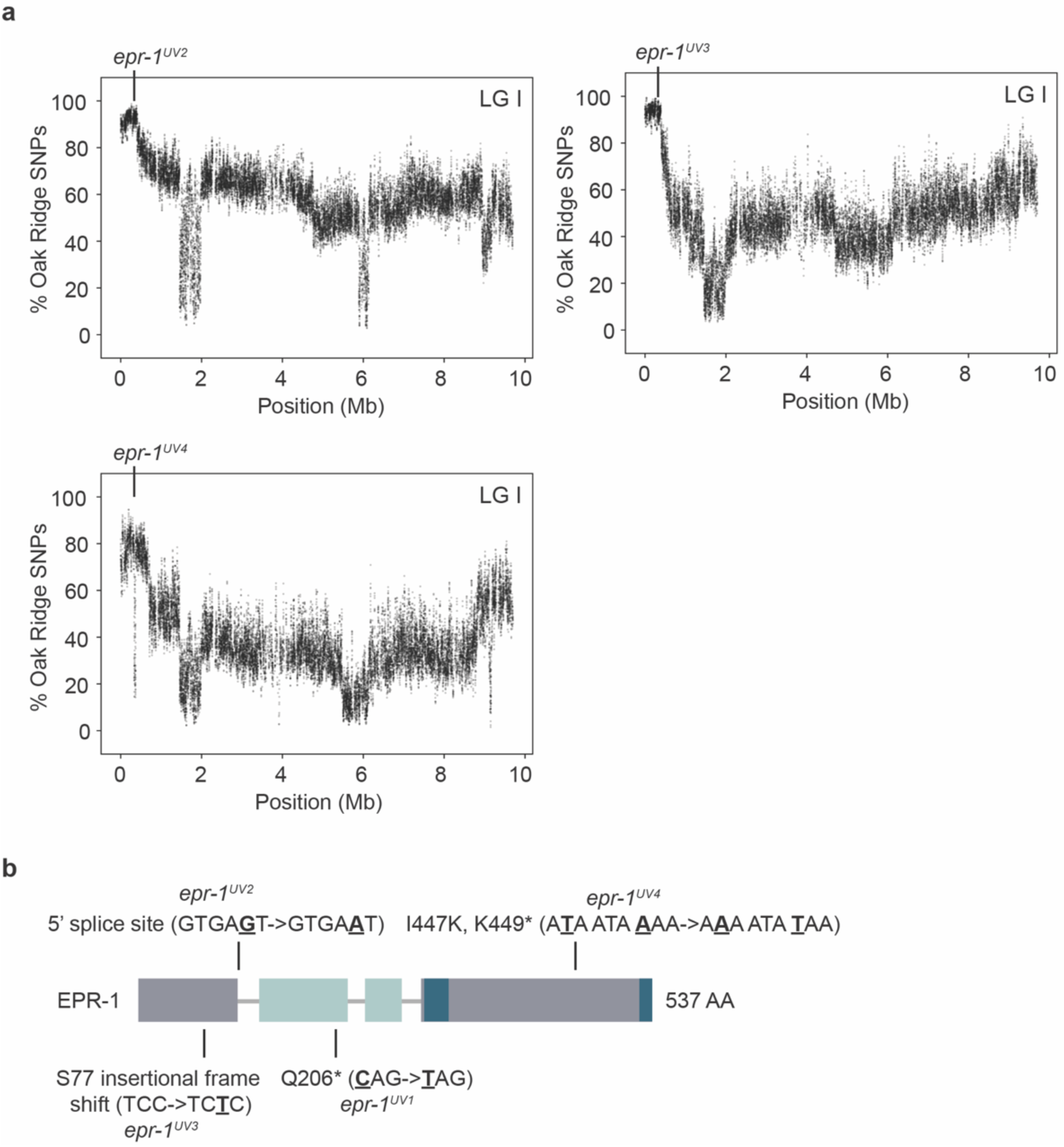
Three additional alleles of *epr-1* were identified during selection for mutants defective in H3K27 methylation-mediated repression. **a**, Whole genome sequencing of pooled genomic DNAs of mutant progeny, isolated from three additional mutants crossed with wild-type Mauriceville, independently identified a region on the left arm of LG I that was enriched for Oak Ridge SNPs and contained mutations in *epr-1* (*NCU07505*). **b**, Domain structure of NCU07505 showing the location and nature of the mutations in the *epr-1* alleles. Exons are indicated as boxes, with the BAH domain in light blue, the PHD finger (split) in dark blue, and exons with no known domains in gray. Gray lines connecting exons indicate the introns.

**Supplementary Fig. 2.**
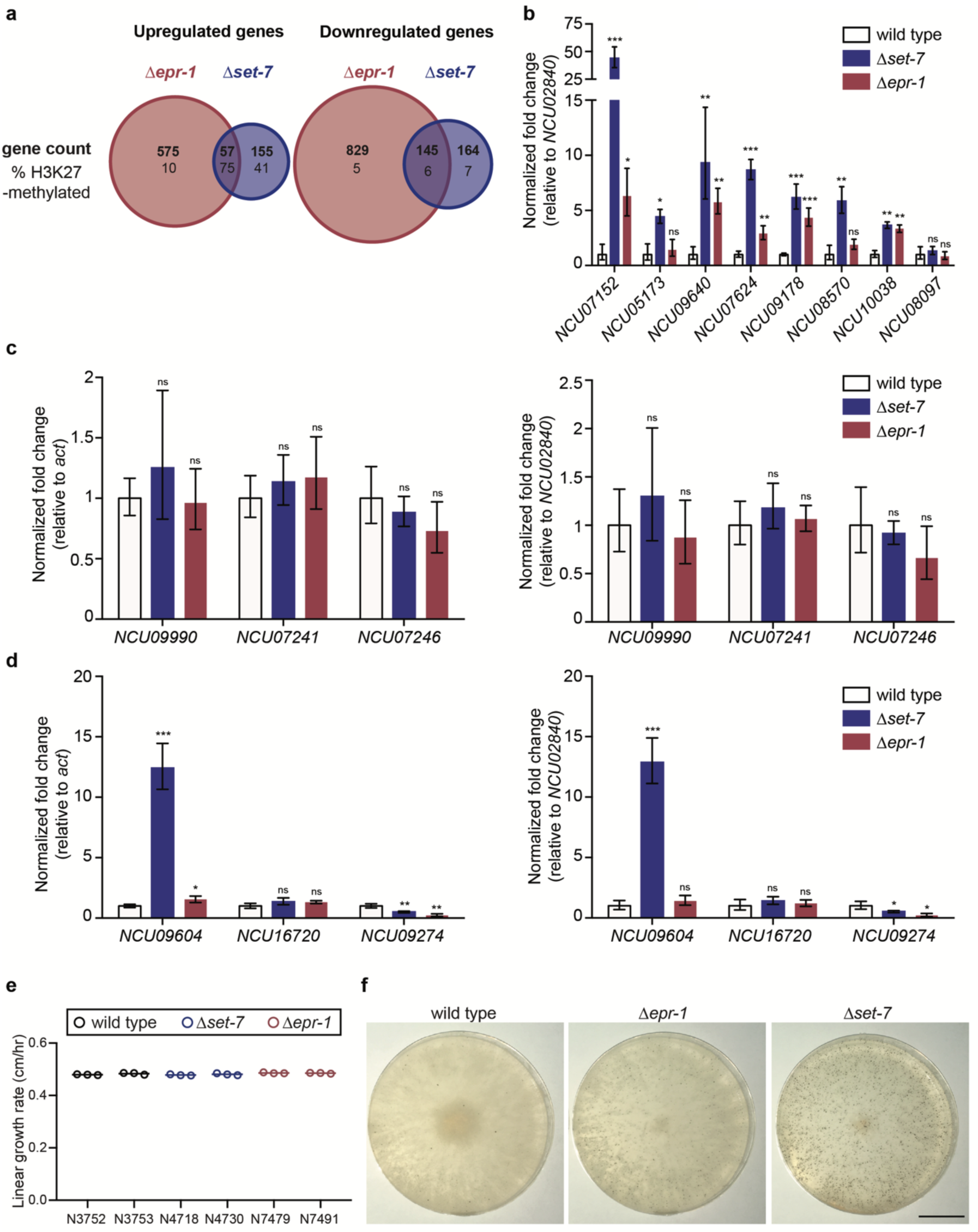
Gene expression and phenotypic analysis of Δ*epr-1* and Δ*set-7* strains. **a**, Bold numbers indicate genes upregulated (left) or downregulated (right) in only *Δepr-1* strains, only *Δset-7* strains, or in both *Δepr-1* and *Δset-7* strains by mRNA-seq. Significance was determined using a cutoff of log_2_(mutant/wild type) > 1 for upregulated genes and < −1 for downregulated genes with a *P* value < 0.05 using the Benjamin-Hochberg correction for multiple comparisons. The percentage of upregulated genes that are marked by H3K27 methylation for each gene set is indicated below the gene count. **b**, RT-qPCR of H3K27-methylated genes that were replaced with antibiotic resistance genes (*NCU07152, NCU05173*), and used for initial selection of mutants, and H3K27-methylated genes that appeared upregulated in both Δ*epr-1* and Δ*set-7* strains by mRNA-seq (*NCU09640, NCU07624, NCU09178, NCU08570, NCU010038, NCU08097*). Each value was normalized to gene expression of *NCU02840* (an alternative housekeeping gene) and presented relative to wild type. **c**, RT-qPCR for three genes (*NCU09990, NCU07241, NCU07246*) that appeared upregulated only in *Δepr-1* strains by mRNA-seq. Each value was normalized to gene expression of *act* (left) or *NCU02840* (right) and presented relative to wild type. **d**, RT-qPCR for three genes (*NCU09604, NCU16720, NCU09274*) that appeared upregulated only in *Δset-7* strains by mRNA-seq. Each value was normalized to gene expression of *act* (left) or *NCU02840* (right) and presented relative to wild type. For all RT-qPCR data, filled bars represent the mean from biological triplicates and error bars show standard deviation (*** for *P* < 0.001, ** for *P* < 0.01, * for *P* < 0.05, and ns for not significant; all relative to wild type by two-tailed, unpaired t-test) **e**, Linear growth rates measured by ‘race tubes’ are shown for two biological replicates of wild-type (N3752, N3753), *Δset-7* (N4718, N4730), and *Δepr-1* (N7497, N7491) strains. Horizontal lines represent the mean of three technical replicates (open circles). **f**, Representative images of biological replicates graphed in Fig. 2c of wild-type, *Δepr-1* and *Δset-7* strains grown unfertilized at 25 °C for two weeks on modified Vogel’s with 0.1% sucrose (31). Overlaid black bar represents 2 cm.

**Supplementary Fig. 3.**
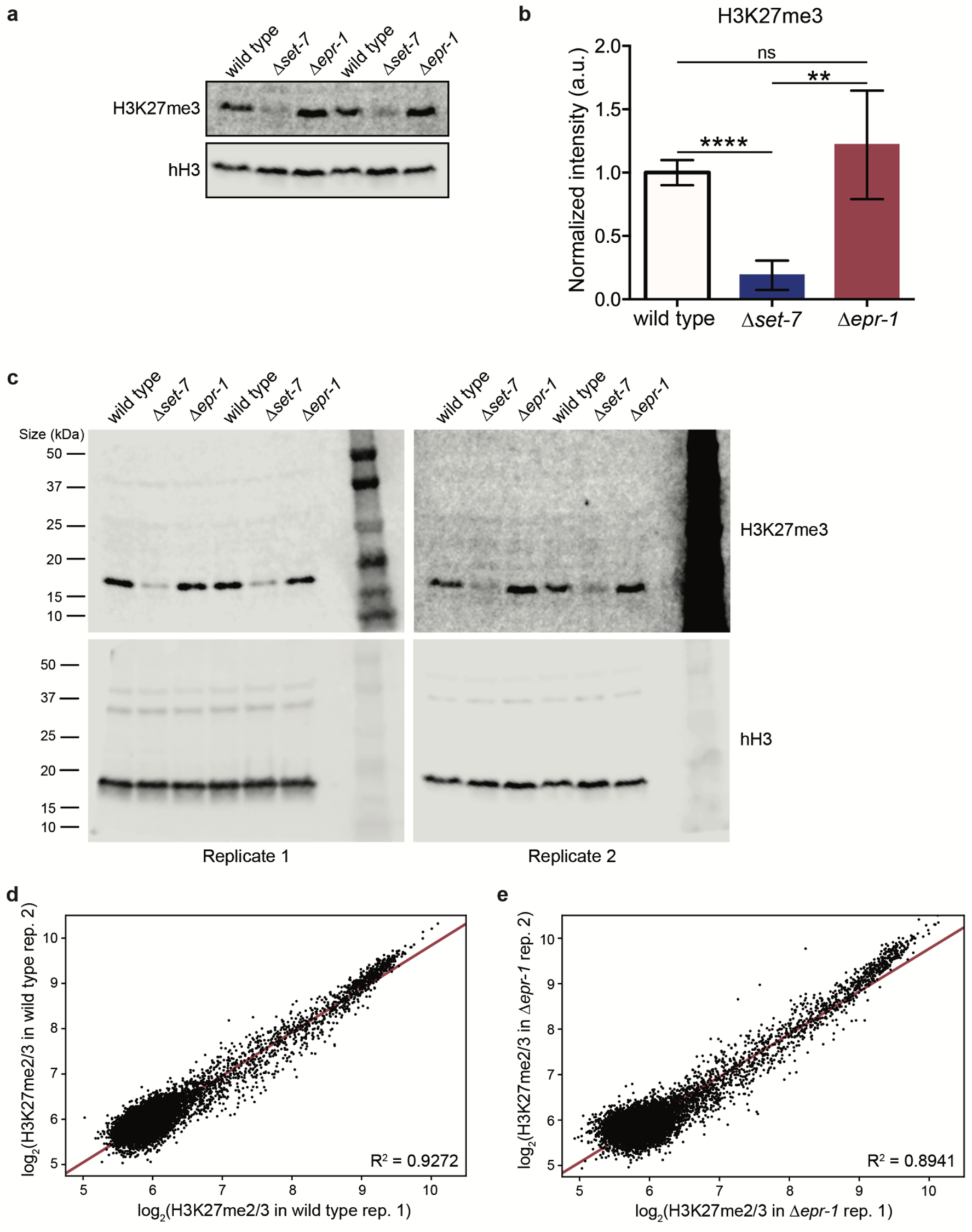
H3K27 methylation levels in *Δepr-1* are comparable to wild type. **a**, Western blot showing H3K27me3 and total histone H3 (hH3) in the indicated strains (additional replicate of experiment illustrated in Fig. 3a). The same lysate was run on separate gels, and hH3 was used as a sample processing control. **b**, Quantification of the H3K27me3 band intensity averaged from 4 biological replicates. Each band intensity is normalized to the corresponding hH3 band and to the wild-type average. Filled bars represent the mean and error bars show standard deviation (a.u. signifies arbitrary units; **** for *P* < 0.0001, ** for *P* < 0.01, and ns for not significant; all relative to wild type by two-tailed, unpaired t-test). **c**, Full blots with molecular weight markers in kilodaltons (kDa) for the cropped images in Fig. 2a and panel **a** (above) are shown. **d**, Scatter plot showing the correlation of H3K27me2/3 levels at all genes (black dots) for the two wild-type ChIP-seq biological replicates. Line of best fit displayed in red (R^2^ = 0.9272). **e**, Scatter plot showing the correlation of H3K27me2/3 levels at all genes (black dots) for the two *Δepr-1* ChIP-seq biological replicates. Line of best fit displayed in red (R^2^ = 0.8941).

**Supplementary Fig. 4.**
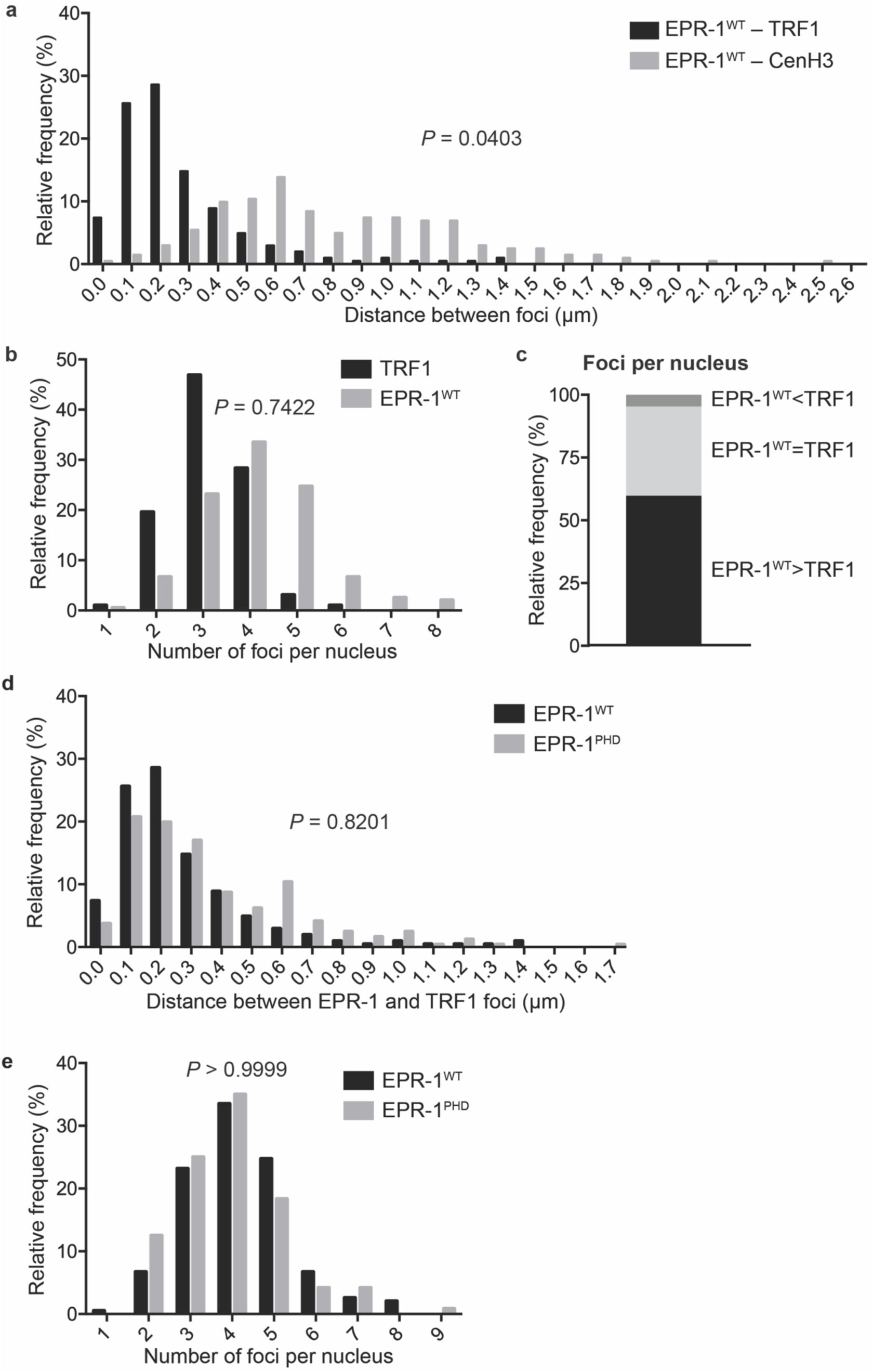
EPR-1^WT^ and EPR-1^PHD^ exhibit no difference in distance to telomeres or EPR-1 foci number. **a**, Histogram shows the relative frequency of distances between an EPR-1^WT^ focus and the closest TRF1 focus (black bars) or CenH3 focus (gray bars) (n = 203, *P* = 0.0403, two-tailed Mann-Whitney test). **b**, Histogram shows the relative frequency of nuclei with the indicated number of TRF1 or EPR-1^WT^ foci (n = 194, *P* = 0.7422, Wilcoxon matched-pairs test). **c**, Stacked bar graph comparing the number of TRF1 and EPR-1^WT^ foci within a single nucleus (n = 194). **d**, Histogram shows the relative frequency of distances between an EPR-1^WT^ focus and the closest TRF1 focus (black bars; n = 203) or an EPR-1^PHD^ focus and the closest TRF1 focus (gray bars; n = 241) (*P* = 0.8201, two-tailed Mann-Whitney test). **e**, Histogram shows the relative frequency of EPR-1 foci per nucleus for EPR-1^WT^ (n = 194) or EPR-1^PHD^ (n = 120) (*P* > 0.9999, Wilcoxon matched-pairs test).

**Supplementary Fig. 5.**
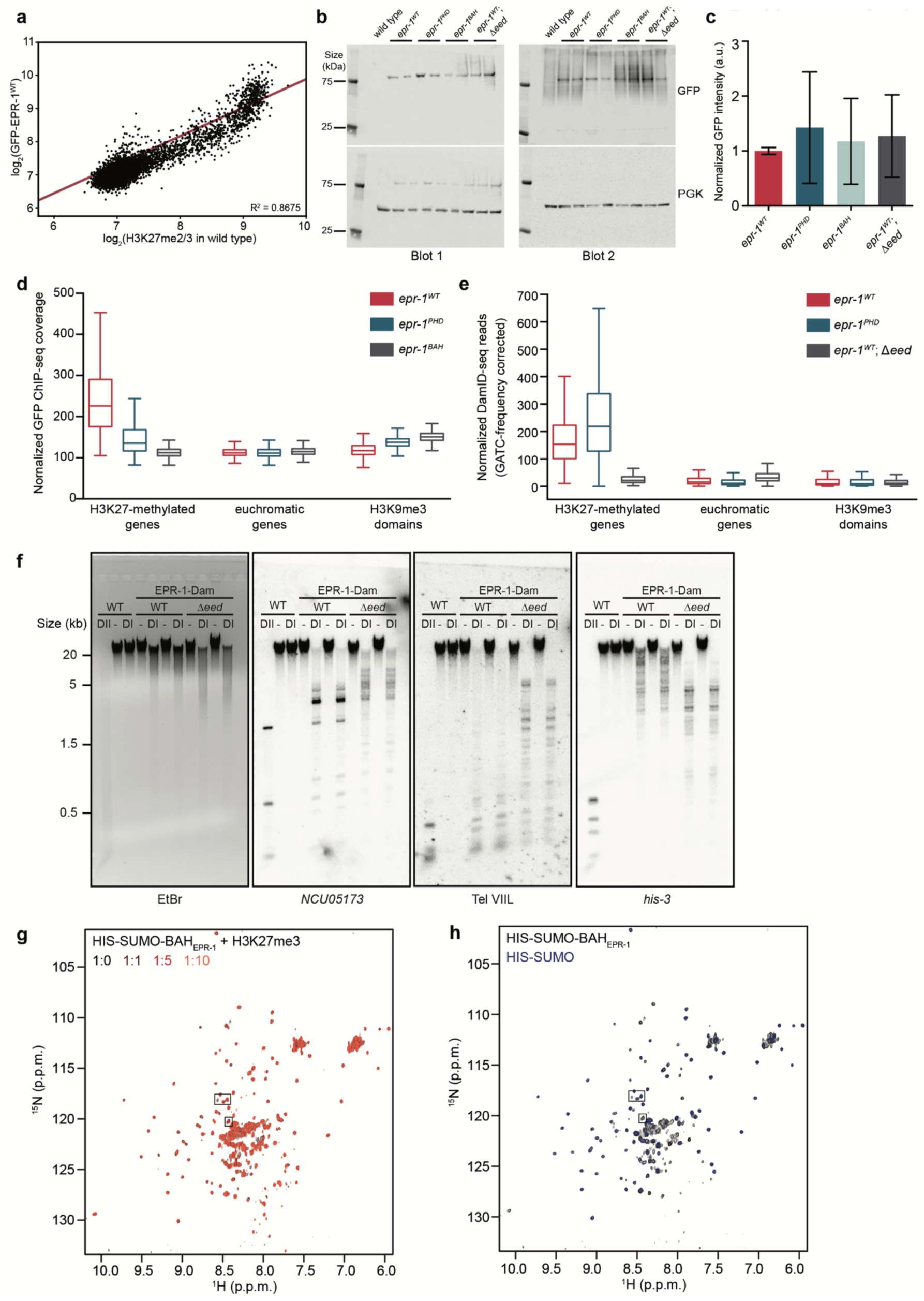
EPR-1 associates with H3K27 methylation *in vivo* and *in vitro*. **a**, Scatter plot showing the correlation of levels of H3K27me2/3 and GFP-EPR-1^WT^ for all genes (black dots), as determined by ChIP-seq. Line of best fit displayed in red (R^2^=0.8675). **b**, Western blot shows GFP-EPR-1 expression in the indicated strains. Phosphoglycerate kinase (PGK) is used as a loading control. Each genotype (except wild type, negative control) was run in biological duplicate and repeated. **c**, Quantification of the GFP band intensity averaged over 4 biological replicates. The intensities are relative to the corresponding PGK band and normalized to the wild-type average from the same blot. Filled bars represent the mean and error bars show standard deviation (a.u. signifies arbitrary units). **d**, Box and whisker plot of normalized GFP-EPR-1 ChIP-seq coverage from *epr-1*^*WT*^, *epr-1*^*PHD*^ and *epr-1*^*BAH*^ strains is shown for the indicated regions of the genome. Box represents interquartile range, horizontal line is median, and whiskers represent minimum and maximum values. **e**, Box and whisker plot of normalized EPR-1 DamID-seq coverage from *epr-1*^*WT*^, *epr-1*^*PHD*^ and *epr-1*^*WT*^; *Δeed* strains is shown for the indicated regions of the genome. Reads have been corrected for the frequency of GATC sites. Box represents interquartile range, horizontal line is median, and whiskers represent minimum and maximum values. **f**, DamID Southern blot of genomic DNA from the indicated strains digested with DpnI (DI), DpnII (DII) or left undigested (-). DpnII, which digests GATC sites without methylated adenines, reveals pattern of complete digestion in wild type. DpnI, which fails to digest GATC sites bearing adenine methylation, reveals the extent of methylation by Dam at probed regions (*NCU05173* and Tel VIIL, H3K27-methylated; *his-3*, euchromatin). Ethidium bromide (EtBr) shows total DNA. Biological replicates are shown. **g**, An overlay of ^1^H-^15^N HSQC spectra of ^15^N-labelled HIS-SUMO-BAH_EPR-1_ fusion in the presence of increasing concentrations of H3K27me3 peptide. **h**, An overlay of ^15^N-labelled HIS-SUMO-BAH_EPR-1_ fusion with the ^15^N-labelled HIS-SUMO alone. Boxed areas are shown enlarged in Fig. 5d. Spectra are color coded as indicated.

**Supplementary Table 1.**
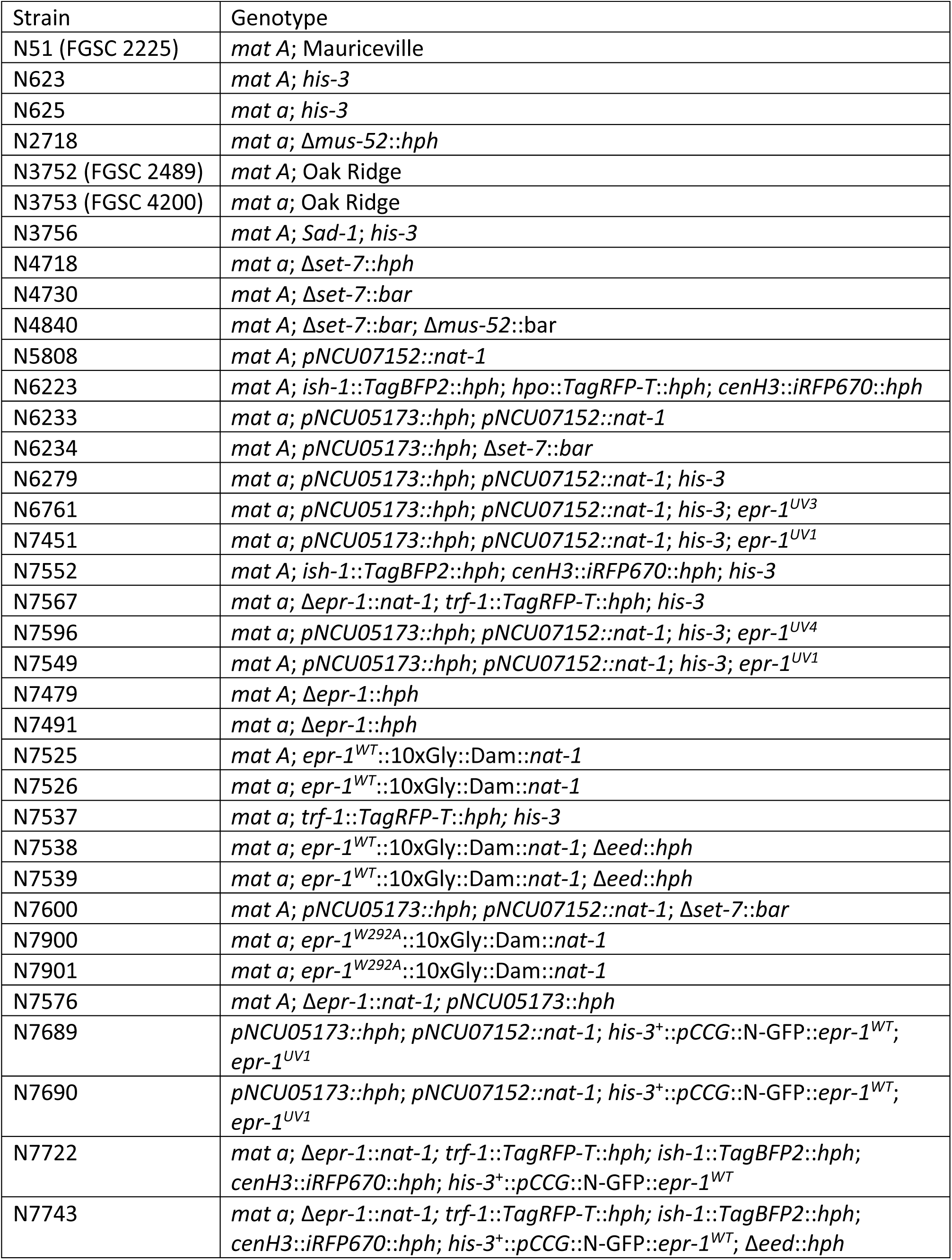
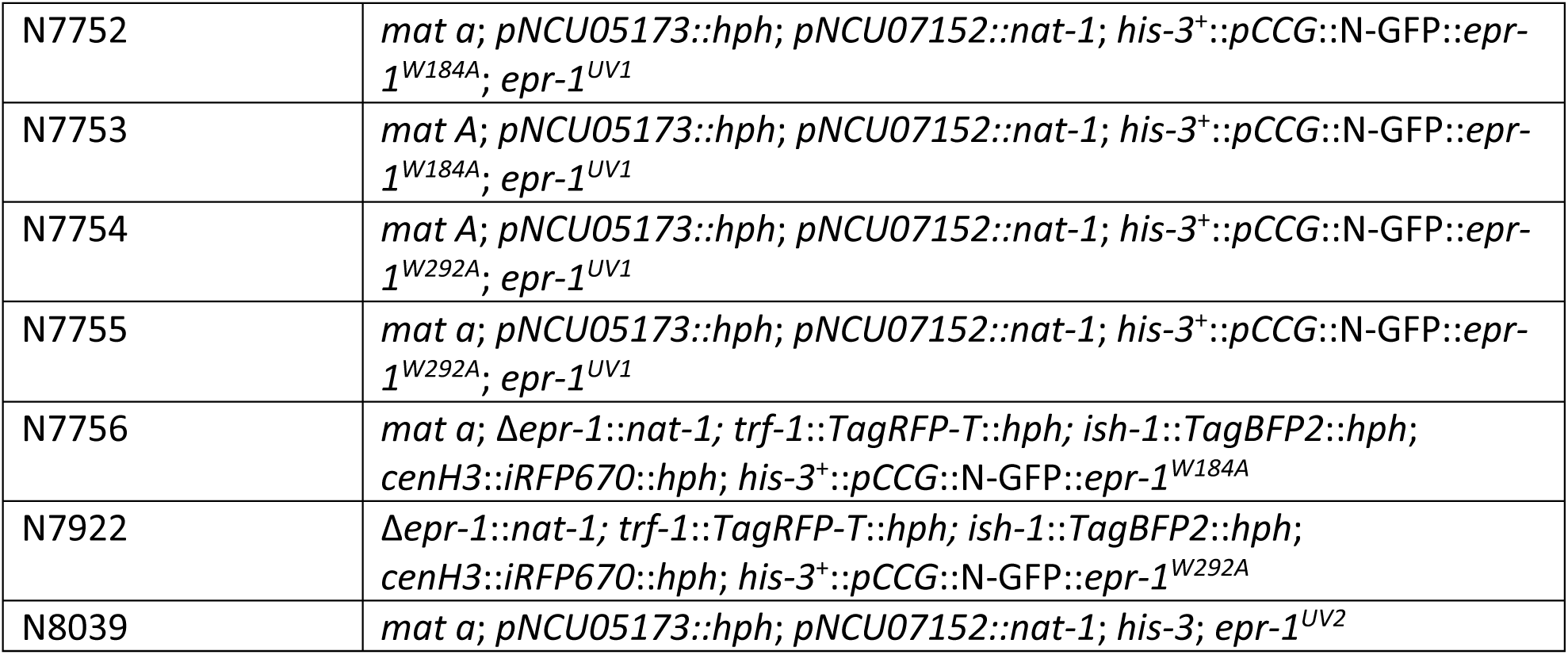
*N. crassa* strains.

**Supplementary Table 2.**
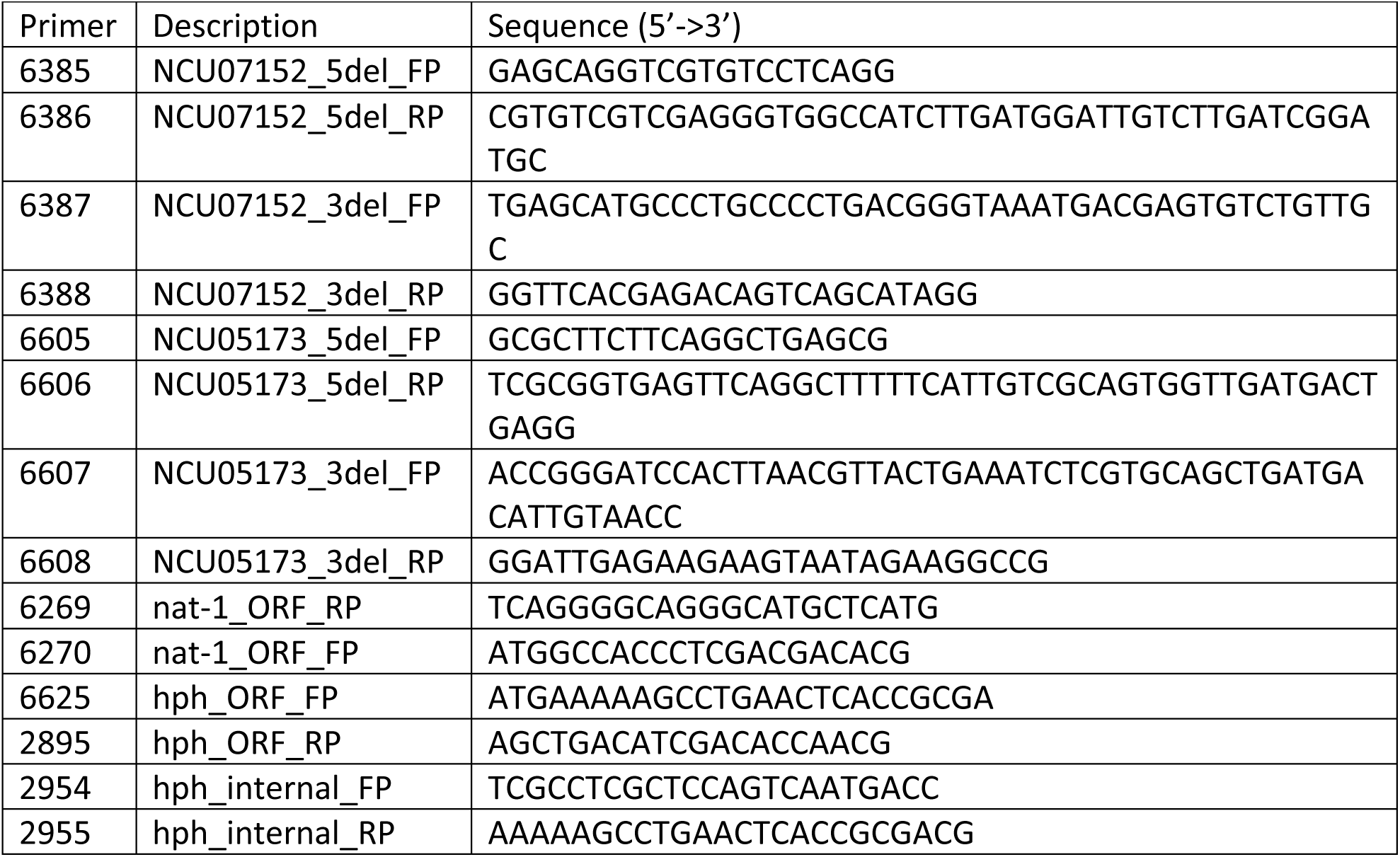
Replacement of NCU05173 and NCU07152 ORFs with *hph* and *nat-1*.

**Supplementary Table 3.**
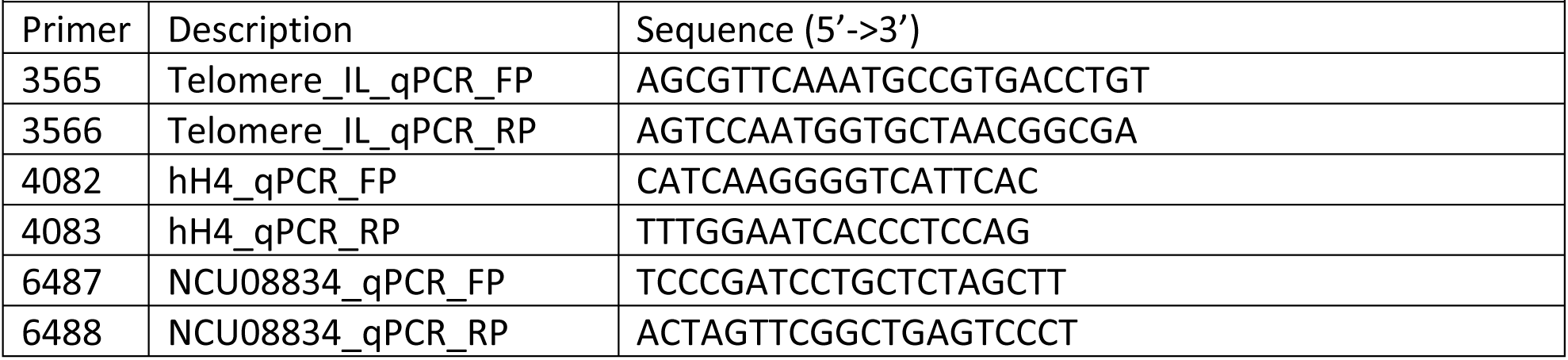
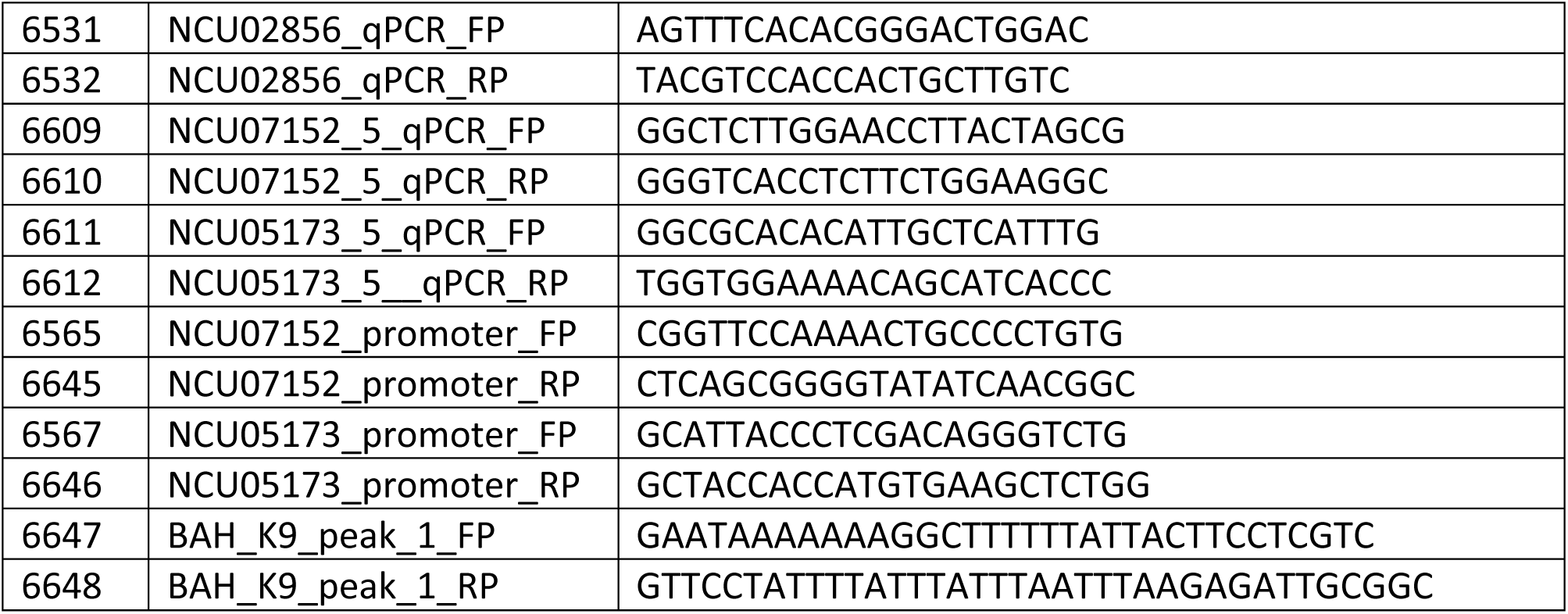
ChIP qPCR primer pairs.

**Supplementary Table 4.**
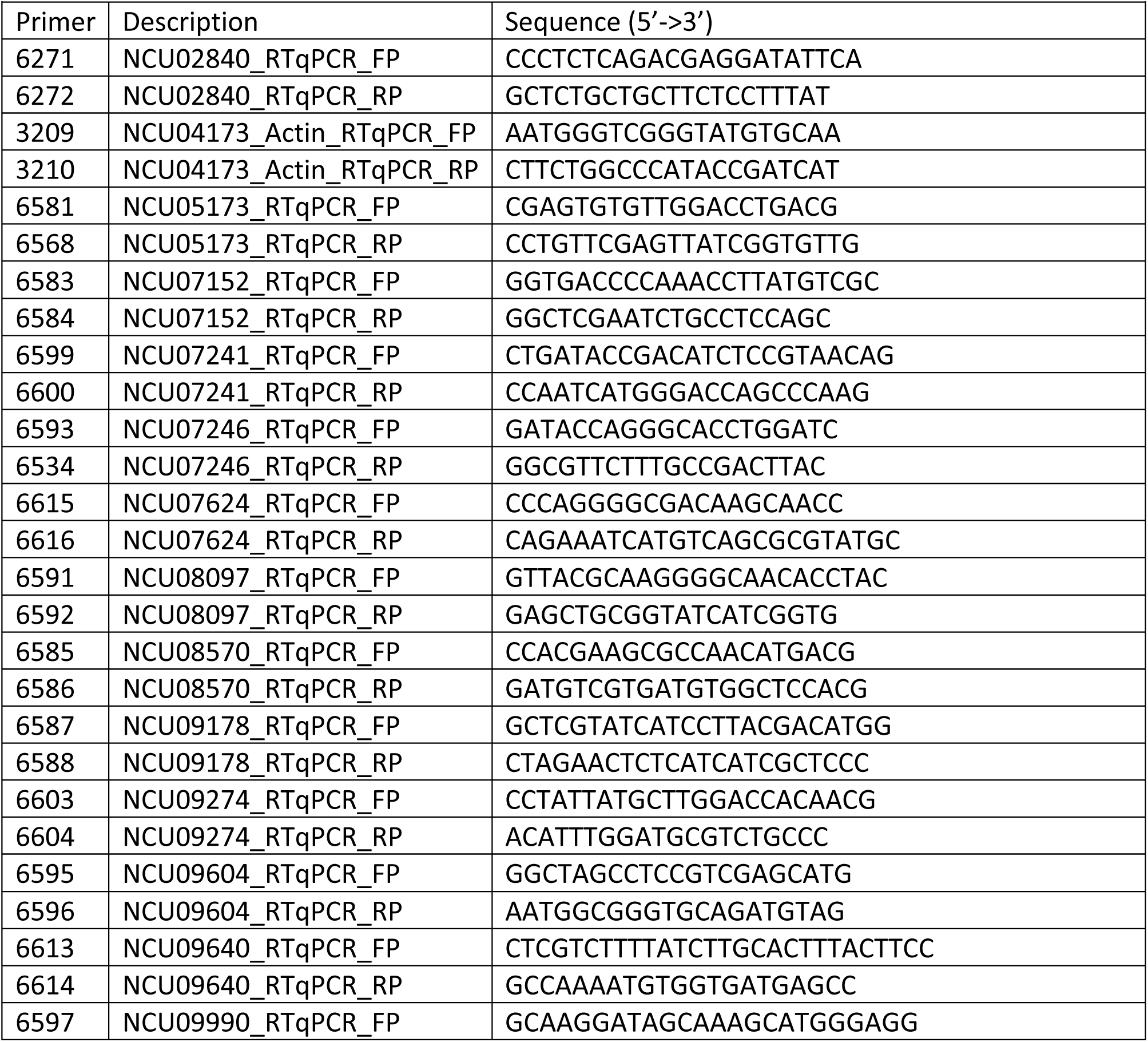
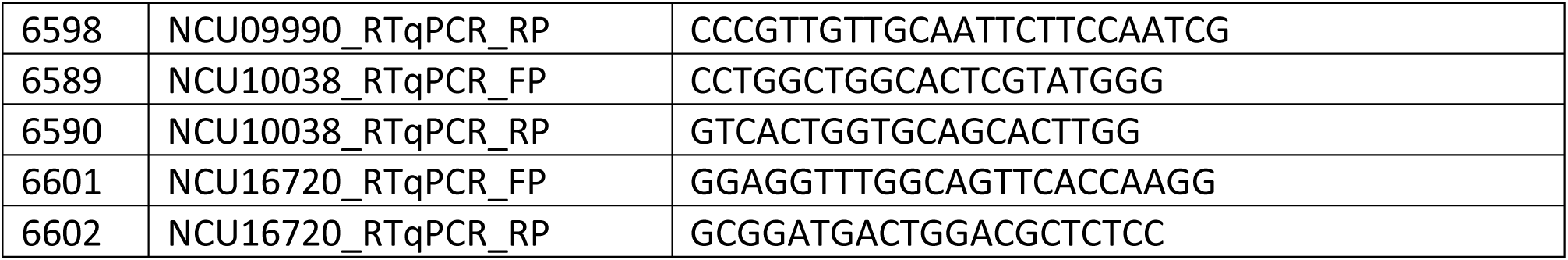
RT-qPCR primer pairs.

**Supplementary Table 5.**
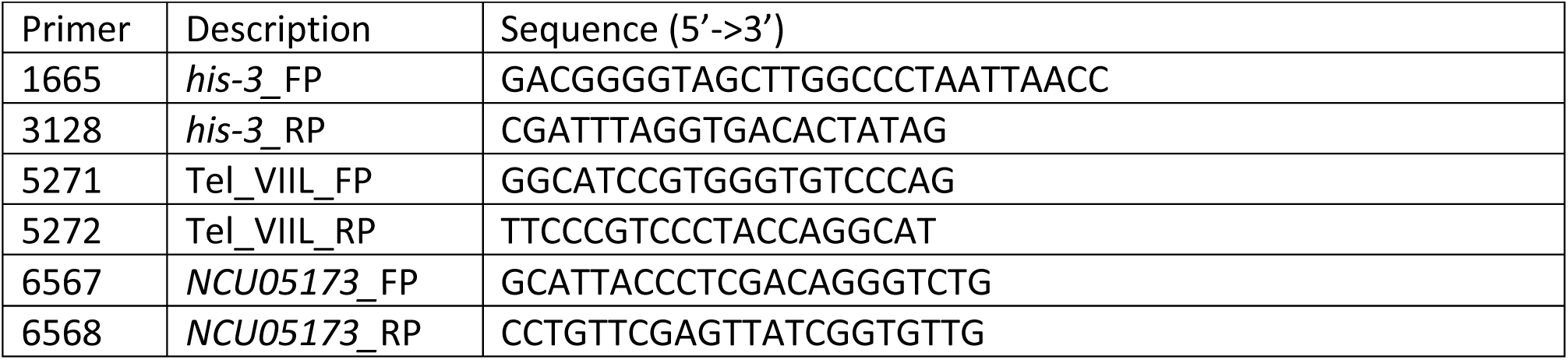
Southern probes.

**Supplementary Table 6.**
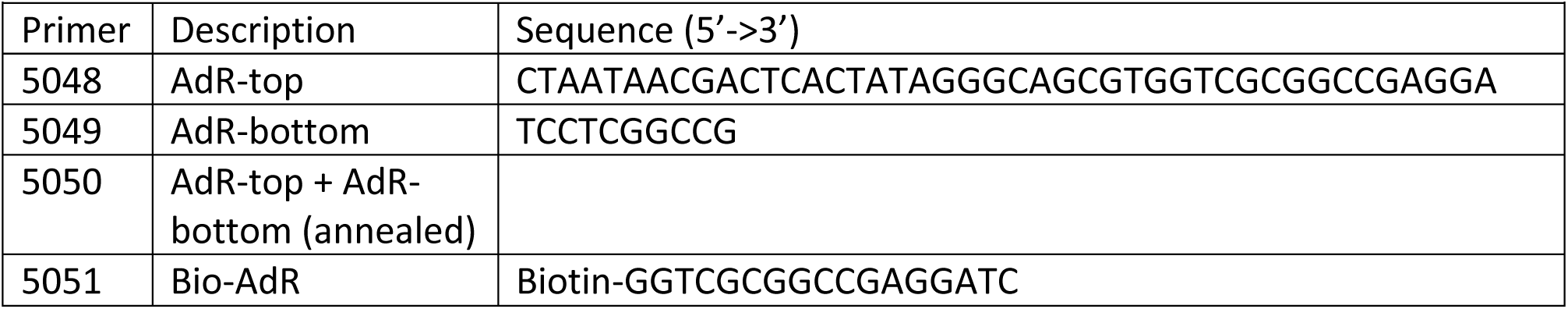
DamID-seq protocol.

**Supplementary Table 7.**
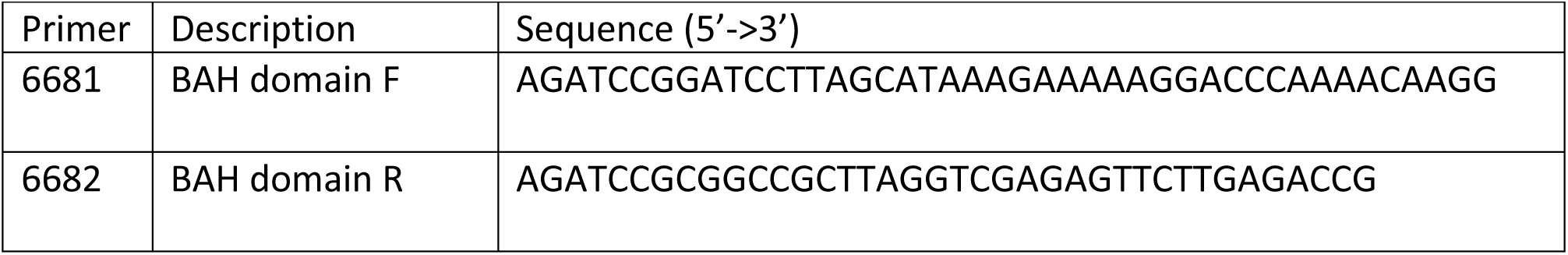
Cloning the BAH domain of EPR-1.

**Supplementary Table 8.** Plasmids.

| Supplementary Table 8. Plasmids |  |
| --- | --- |
| Plasmid | Description |
| 1991 | pBM61 – *his-3*<sup>+</sup> targeting vector |
| 2406 | *his-3*<sup>+</sup>::*pCCG*::N-GFP |
| 3329 | *his-3*<sup>+</sup>::*pCCG*::N-GFP::*NCU07505* |
| 3345 | *his-3*<sup>+</sup>::*pCCG*::N-GFP::*NCU07505*<sup>W184A</sup> |
| 3346 | *his-3*<sup>+</sup>::*pCCG*::N-GFP::*NCU07505*<sup>W292A</sup> |
| 3130 | pZero::10xGly::3xFLAG::*trpC*::*nat-1* |
| 3131 | pZero::10xGly::Dam::*trpC*::*nat-1* |
| 3237 (FGSC 10598) | pAL12 – source of *nat-1* |
| 2283 | pCSN44 – source of *hph* |
| 3230 | pE-SUMO |
| 3410 | pE-SUMO + BAH<sub>EPR-1</sub> |

**Supplementary Table 9.**
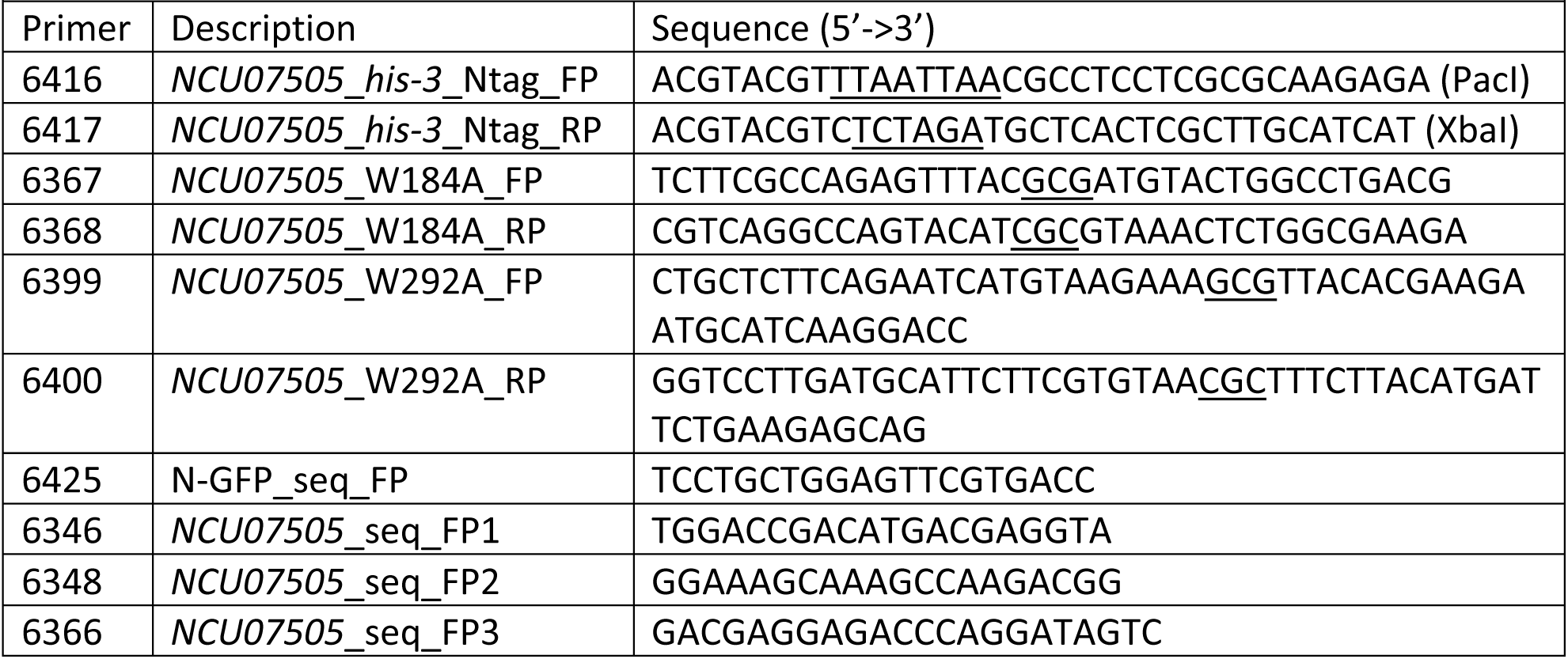
Generation and verification of *his-3*^+^::*pCCG*::N-GFP::EPR-1 constructs.

**Supplementary Table 10.**
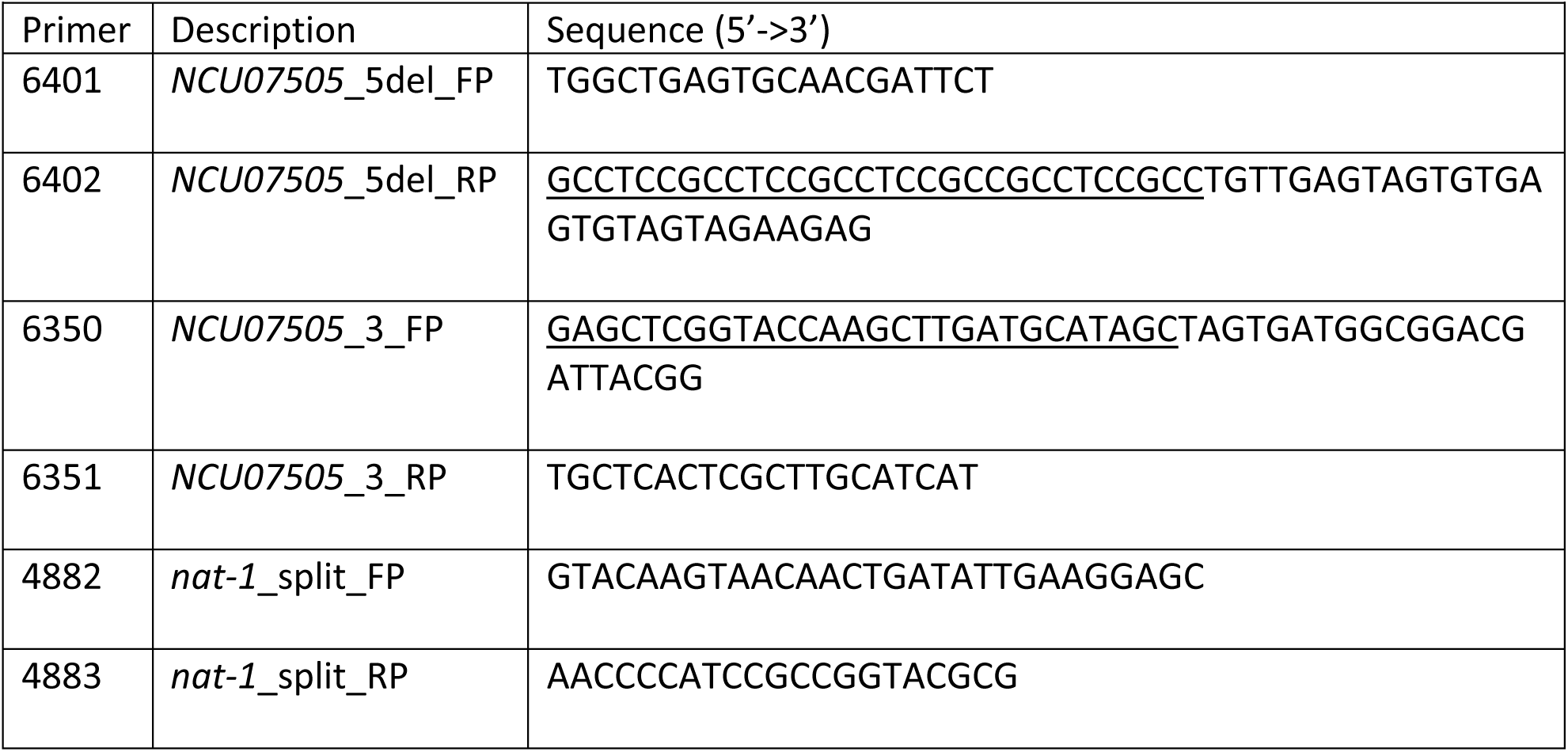
Replacement of *epr-1* with *trpC::nat-1*.

**Supplementary Table 11.**
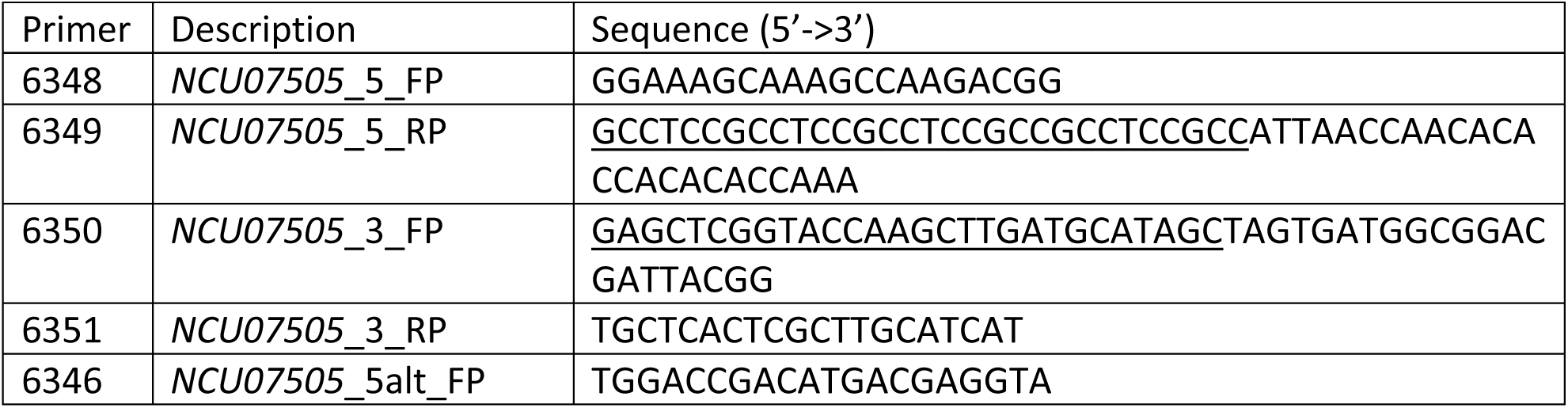
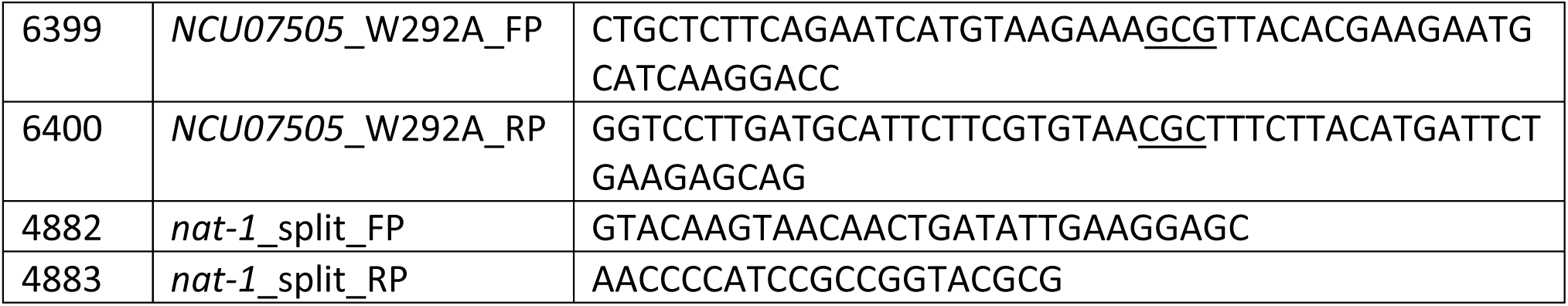
Endogenous tagging of EPR-1 with 10xGly::Dam (WT and PHD)

